# Balanced spatiotemporal color responses are fine-tuned to natural light spectrum in mice ventral retina

**DOI:** 10.1101/2025.03.13.643035

**Authors:** Tom Quétu, Awen Louboutin, Filippo Castellani, Remi Baroux, Ulisse Ferrari, Matías A. Goldin

**Affiliations:** Sorbonne Université, CNRS, Inserm, Institut de la Vision, F-75012 Paris, France; Politecnico di Milano, Department of Electronics, Information, and Bioengineering, Milan, Italy; The BioRobotics Institute, Scuola Superiore Sant’Anna, Pisa, Italy

## Abstract

Color vision is vital for animal survival, essential for foraging and predator detection. In mice, as in other mammals, color vision originates in the retina, where photoreceptor signals are processed by neural circuits. However, retinal responses to stimuli involving multiple colors are still not well understood. One possible explanation of this knowledge gap is that previous studies have not thoroughly examined how neuronal activity adapts to a 30 seconds to a few minutes timescale when exposed to multiple color sources. To address this, we systematically varied the UV-to-green light balance with a custom-built stimulator targeting mice opsins spectra while recording retinal ganglion cell responses across the dorso-ventral axis of the retina using multielectrode arrays. Responses to full-field chirp and checkerboard stimulations with alternating UV and green light revealed that more than one order of magnitude of intensity difference favoring green M-opsin over UV S-opsin is needed for a balanced reliability in retinal ganglion cell responses in the ventral retina. An incorrect balance, with slightly increased UV light, silenced responses to green illumination. To determine if these values are consistent with natural conditions, we analyzed isomerisation rates in the mouse retina across different times of the day. We found that the M- to S-opsin activation ratio remains constant through the mesopic-photopic range, and that our empirically determined values in the ventral retina align well with these natural conditions. These lie far from a simple equalization of M- and S-opsin isomerisation rates, which we found only balances ganglion cell responses in the dorsal retina. In conclusion, a finely tuned color intensity balance matching natural light spectrum is essential for accurately measuring both fast temporal responses and detailed spatial receptive fields in the ventral retina.

## 1. Introduction

The number, the type and the distribution of photoreceptors across the retina vary widely among vertebrates, reflecting evolutionary adaptations to their behavior. Radically different configurations can be observed in diurnal and nocturnal animals, as well as depending on their lifestyle, such as whether they are prey or predators (Baden, 2024; Baden, Euler & Berens, 2020; Peichl, 2005; Peichl et al., 2019; Pessoa et al., 2014; Yoshimatsu, 2020). These adaptations optimize the visual information they can obtain in natural environments, directly influencing their ability to find food, detect predators, and navigate. Color vision, which plays a key role, relies on comparing signals from photoreceptors with different spectral sensitivities determined by their opsins (Osorio et al., 2004; Valenta et al., 2013).

The mouse retina, like in many other mammals but primates (Dacey, 1996; Dacey, 1999), expresses two types of cone opsins, S-Opsin and M-Opsin, with peak sensitivities at 360 nm and 508 nm respectively (Nikonov et al., 2006). Unlike primates, robust cone-opponent cells, crucial for color discrimination, have not been clearly identified in the mouse retina, with the exception of one intrinsically sensitive ganglion cell (Stabio et al., 2018). In primates, these cells compare input from different types of cones, enabling wavelength discrimination (Field et al., 2010). Some studies suggest that such circuits might coexist in the thalamic region of mice (Mouland et al., 2021), while others propose that certain retinal ganglion cells use rod-cone opponency for color discrimination (Joesch & Meister, 2016; Khani & Gollisch, 2021). Recent studies investigated the origin of color opponency in the ventral retina, tracing it back to interactions at the cone synapse through lateral inhibition from horizontal cells, partially driven by rods (Behrens et al., 2016; Yoshimatsu et al., 2021). Furthermore, bipolar cells relay the chromatic information to retinal ganglion cells, where type-specific, nonlinear center-surround interactions result in color-opponent output channels to the brain (Szatko et al., 2020; Korympidou et al., 2024). Behavioral studies confirm that mice can distinguish between different colors, particularly in the upper visual field, likely due to the distribution of photoreceptors in the retina (Denman et al., 2018).

The expression patterns in the mice retina of the two color opsins create opposing and overlapping gradients along the dorsal-ventral axis (**Fig. 1a**) (Szel et al., 1992). As a result, the majority of cones express both opsins. However, there are also cones that exclusively express one opsin. Thus, S-Opsin enrichment in the ventral retina may help the detection of short-wavelength light from the sky, and M-opsin in the dorsal retina permits the perception of the ground (e.g., a grassy field), while co-expression of both opsins broadens the spectral range of individual cones and improves perception under varying conditions of ambient light. A recent study suggested that the nonlinear interaction of UV and green would serve to delineate the threshold between the sky and the ground (Khani & Gollisch, 2021). The ventral portion of the mice retina emerges as an ideal system for investigating color processing capabilities (**Fig. 1b**). Moreover, although the role of rods in color discrimination is not fully understood, there is a close overlap in spectral sensitivity with the M-opsin (Wang, Weick & Demb, 2011; Govardovskii et al., 2000). Their influence on the overall retinal response is significant, particularly in their potential contribution to green color detection at high illumination levels (photopic regime) (Adelson, 1982; Khani & Gollisch, 2021; Tikidji-Hamburyan et al., 2017; Pearson et al., 2015).

**Figure 1.**
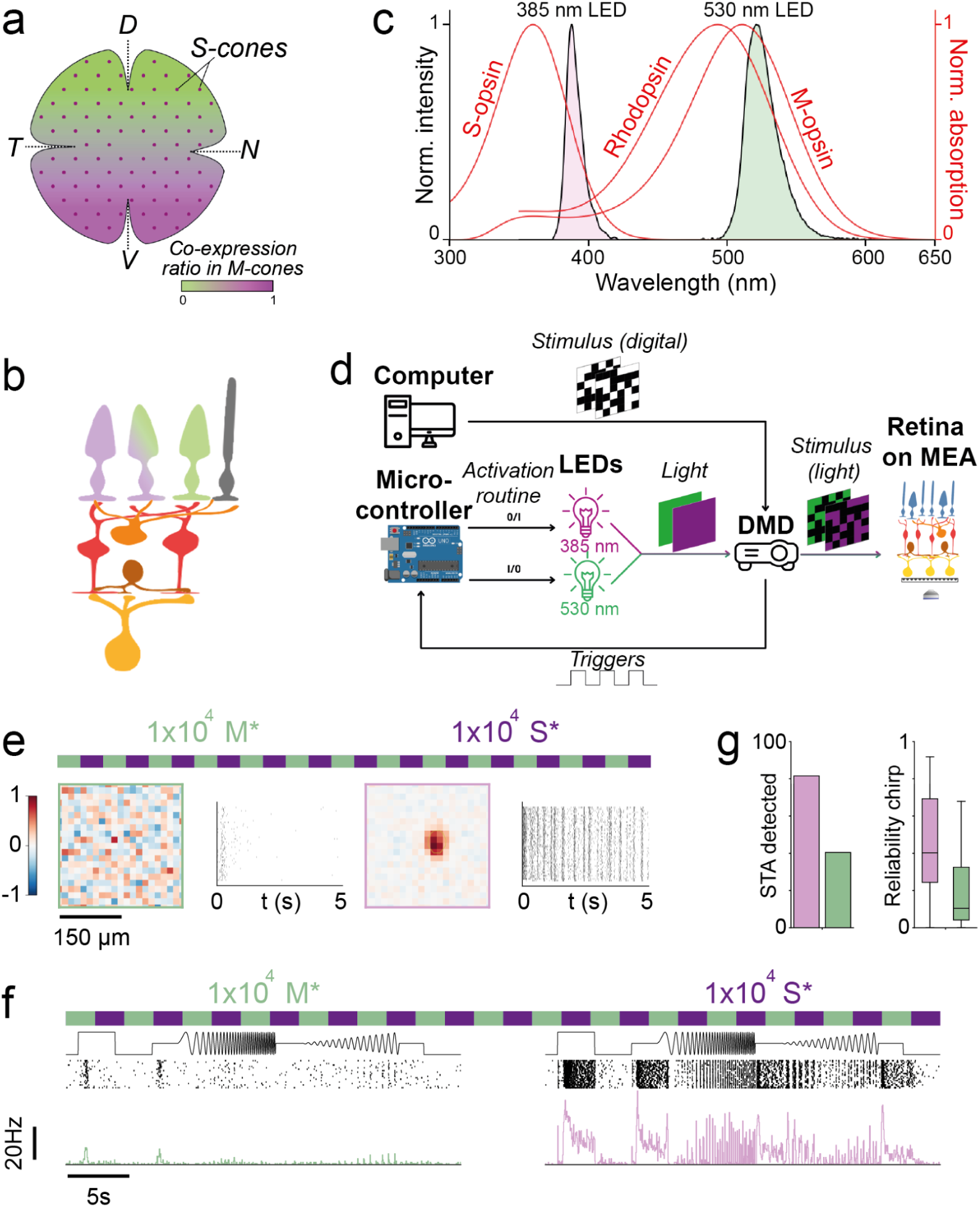
Interleaved identical activation of S- and M-opsin prevents high resolution and high frequency ganglion cells’ responses to green light in the ventral retina. **a.** Schematic distribution of cone photoreceptors across the mouse retina. Dots represent distribution of true S-cones and shading co-expression ratio of S- and M-opsin in M-cone. d: dorsal, n: nasal, v: ventral, t: temporal. **b.** Schematic of the information pathways from photoreceptors to a ganglion cell. Top: Green (M-Opsin), UV (S-Opsin), Green+UV (M-S-opsin co-expression) and Gray (rod). Information is relayed to and processed by middle layer cells (horizontal, bipolar and amacrine) before reaching a ganglion cell (bottom). **c.** The normalized absorption spectra of S-Opsin, M-Opsin and rhodopsin overlaid with the normalized power spectra of UV (385 nm) and green (530 nm) LEDs. The opsins have overlapping absorption spectra, with M-Opsin and rhodopsin extending into shorter wavelengths. This overlap makes it impossible to stimulate S-Opsin without also affecting M-Opsin with the 385 nm LED, though the effect is small. **d.** Schematic of the experimental setup for Multi-Electrode Array (MEA) recordings with a color stimulation device. Two LEDs (385 nm and 530 nm) are modulated via a microcontroller, combined into a single beam using dichroic mirrors, and transmitted through an optical fiber to a Digital Mirror Device (DMD). The light is then focused onto the retina, positioned with the ganglion cell layer in contact with the MEA electrodes, on the back focal plane of a microscope. **e.** Example cell STA and spike raster responses to an interleaved green (left) and UV (right) checkerboard (40 s green checkerboards alternated with 40 s Violet checkerboards). The horizontal green and violet bars represent the interleaved color stimuli. Scale bar: from dark (blue) to bright (red). **f.** Example cell PSTH and spike raster response to an interleaved green (left) and UV (right) chirp (32s bouts). **g.** Quantification of the number of detected STAs and reliability of the chirp responses in the illumination configuration corresponding to panels e and f.

Understanding retinal visual processing has long relied on simplified, artificial stimuli, which have been instrumental in uncovering key properties of the retina (center-surround antagonism, non-linear spatial integration, contrast computation, etc.). However, recent studies using natural images and videos (Höfling et al., 2024; Franke et al., 2022) emphasize the importance of natural scene statistics (Tkačik et al, 2011; Atick et al., 1992; Doi et al., 2003) in revealing complex nonlinear processes within the retinal circuit (Goldin, Lefebvre, Virgili et al., 2022; Goldin et al., 2023; Goethals et al., 2025; Karamanlis et al., 2022; Turner et al., 2018; McIntosh et al., 2016; Maheswaranathan et al., 2023). While artificial stimuli remain valuable for probing specific retinal functions, they do not fully capture how the retina operates in real-world conditions, making natural stimulation a fundamental tool. The incorporation of color into these stimuli can significantly enhance our knowledge of retinal circuits (Khani & Gollisch, 2021; Höfling et al., 2024; Qiu et al., 2021; Szatko et al., 2020; Fornetto et al., 2024). However, despite technological advancements that allow for precise measurement of color information in natural scenes (Franke et al., 2019; Qiu et al., 2021), there are still limitations on the length of experiments where these stimuli can be presented in vitro. This poses challenges, especially when investigating the nonlinear interactions that occur when multiple colors are present.

Natural scenarios can have different color content, such as night or daylight illumination. Therefore, depending on the animal adaptation to its environment, the ratio between the intensities of the chromatic information present in a natural stimulation can have different impacts on the nonlinear processing of visual information by the retinal circuit. In addition, light entering the eye can be differentially absorbed by the optics media of the eye depending on the wavelength of the incident light, thus modifying the chromatic content of natural scenes arriving at the retina (Henrikson et al., 2010; Douglas & Jeffery, 2014). Shorter wavelengths will be less transmitted, thus having less intensity at the photoreceptor layer of the retina at the back of the eye. Thus, in order to approach a more naturalistic stimulus design, the natural illumination spectrum should be taken into account.

Studying color processing in the ex-vivo retina can also be impacted by the recording technology chosen. Considerable light activation of the photoreceptors can occur when using functional imaging to record retinal activity. Single or 2-photon microscopy aimed at activating different types of indicators, such as Calcium or glutamate indicators, can put the retina in a high mesopic/low photopic light regime (Euler, Franke & Baden, 2019). The chromatic content of the activation light is not flat in the light spectrum but is considerably higher where the M-opsin presents its higher absorption. This issue has been recognized and its effect is accounted as a contrast reduction of the stimulus projected to the retina. But, can the chromatic unbalance of this background have other effects on retinal processing?

Our study here thus aims to explore the effect of changing the ratio of S-opsin to M-opsin isomerisation rates on retinal responses when using color stimuli. Our goal is to identify the UV and green light intensities that best allow us to study retinal computations involving both colors in mice. To guide our search, we will assess the reliability of the responses of ganglion cells for different color configurations and quantify their responses to fast temporal and fine spatial features that we will test using a chirp and checkerboard stimuli respectively. In the following sections, we will first detail our methods, including the experimental setup and the versatile color stimulation tool we developed, along with the selected stimuli used to test retinal responses. We will then present our results, highlighting the S-, M-opsin isomerisation rates ratios that best balance the responses of ganglion cells in the ventral and dorsal retina. We will finally compare our empirically found ratio to the ones found in a natural setting at different times of the day. Finally, we will discuss the implications of our findings.

## 2. Methods

### 2.1 Electrophysiological recordings

Electrophysiological data were obtained from isolated retinas of eight female C57BL6J mice, aged 4 to 17 weeks. The experiments adhered to the EU Directive 2010/63/EU on the protection of animals used for scientific purposes. After euthanizing the animal, the eye was enucleated and swiftly transferred to oxygenated RINGER medium (for 2L: NaCl 125 mmol/L (14.61 g), KCl 2.5 mmol/L (0.373 g), MgCl₂·6H₂O 1 mmol/L (0.407 g), NaHCO₃ 26 mmol/L (4.369 g), CaCl₂ 1 mmol/L (0.294 g), L-glutamine 0.43 mmol/L (0.126 g), NaH₂PO₄ 1.25 mmol/L (0.300 g), Glucose 20 mmol/L (7.206 g)). Dissection was carried out under dim red light. A piece of the retina (3-5 mm x 3-5 mm) was mounted onto a transparent dialysis membrane and positioned with the ganglion cell side facing a 252-channel multi-electrode array (MEA) with electrodes of 8µm of diameter and spaced 30 μm apart (450 μm x 450 μm). Throughout dissection and recording, the tissue was perfused with oxygenated RINGER solution using a peristaltic perfusion system with two independent pumps: PPS2 (Multichannel Systems GmbH). Retinas were maintained at 36 degrees during the entire experiment.

Electrical activity was extracted from the MEA with an amplifier (Multi Channel System). This data was sampled at 20 kHz and visualized with a dedicated software (MC Rack, Multi Channel System) then exported for analysis (MC Data Tool, Multi Channel System). The raw signals were high-pass filtered at 100 Hz, and spikes were detected using SpyKING CIRCUS (Yger et al., 2018). Subsequent analysis was performed using custom Python scripts. We extracted the activity of 1249 neurons, retaining only those cells with a low number of refractory period violations (< 0.5%, with a median at 0.052% across 7 experiments, 2 ms refractory period) and well-distinguished template waveforms. For the typing experiment, we selected 217 neurons with a slightly higher refractory period violation to aid the clustering procedure (< 1.5%, with a median at 0.172%, 2 ms refractory period) and well-distinguished template waveforms. These criteria ensured the high quality of the reconstructed spike trains.

### 2.2 Color stimulation

Two LEDs (Thorlabs Inc.) with peak wavelengths of 385 nm and 530 nm were used as light sources (**Fig. 1c**). Their intensity was adjusted via a microcontroller (Arduino Uno) connected to a multi-channel 12-bit Digital-to-Analog Converter (Module PmodDA4, DILIGENT). The output from the converter was routed to a custom-built constant voltage multiplier for each LED, allowing an output range of 0-5V. Through SPI communication with a computer, we varied the analog voltage values on individual channels. These values were input to a voltage-to-light intensity controller (Thorlabs Inc.) with a maximum frequency of 1 kHz. The light beams from the LEDs were combined into a single multicolor beam using dichroic mirrors and then directed through an optical fiber to a Digital Mirror Device (DLP9500, Texas Instruments). The pixel size obtained at the retina is 3.5 μm. This color stimulation tool has the capability of changing light intensity value and color at a high frame rate. The combined light was focused on the photoreceptors using standard optics and an inverted microscope (Nikon). The schematic of the setup is shown in **Fig. 1d**.

### 2.3 Light intensity measurements

We measured the light intensity curves *P*(*λ*) of the LEDs, where *λ* denotes the wavelength, using a high resolution spectrometer (USB 200+, Ocean Optics). The total power across the LED spectrum was measured with higher absolute sensitivity using a power meter compact photodiode (PM100D, Thorlabs). Neutral density reflective filters of 15 OD (Thorlabs) were used to decrease the power of the LEDs during the experiments, in order to remain in the central dynamic range of the LED voltage controller when changing the intensity by several orders of magnitude.

### 2.4 Isomerisation rates calculation

The continuous theoretical form to compute the photoisomerisation rate *N* ([isomerisation events/(photoreceptor . second], which we will note from now on [iso/ph.s]), given an emission spectrum *P*(*λ*) in [photons/sec·µm²·nm] is expressed as (Breuninger et al., 2011):

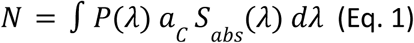

in discrete terms, this formula becomes:

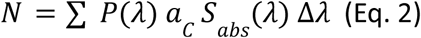

where *P*(*λ*) is the stimulation light spectrum, *S_*abs*_* is the normalized absorption spectrum for the photoreceptor’s opsin with an amplitude equal to 1 at the peak wavelength and *a*_*C*_ is the “effective light collecting area” of the opsin measured in µm². This effective light collecting area takes into account several factors affecting the probability of photon absorption by opsins: quantum efficiency, which depends on the wavelength, the concentration and availability of opsins (i.e., transmembrane proteins not already engaged in the photoisomerisation cycle) and geometric factors of the photoreceptor. For mice, *a*_*C*_ has been experimentally determined to be 0.2 µm² for cones (Breuninger et al., 2011) and 0.5 µm² for rods (Nikonov et al., 2006).

The absorption spectrum *S*_*abs*_ is specific to each opsin and can be sourced from existing literature for cones (**Fig. 1c**, Stockman & Sharpe, 2000). For the rhodopsin curve we used the curve from (Rieke-Lab /calibration-resources github repository), whose short wavelength spectrum lies in between the curve from (Stockman & Sharpe, 2000) and the universal template from (Govardovskii et al., 2000). The remaining values needed to compute isomerisation rates come from the light spectra of each LED *P*_1_ (*λ*), *P*_2_ (*λ*). We measured them after they have traversed the transparent MEA, to account for reflections and optical losses. This is important if the lenses used for the light imaging path are coated for the visible spectrum with a cutoff around 400 nm. These light spectra were discretized with a step size of Δ*λ*=0.5 nm.

With this data we could calculate the isomerisation rate for each photoreceptor’s opsin by summing the effect of each LED:

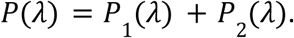

By adjusting the intensity of each LED, different balance between the two LED spectra can be created using the following formula:

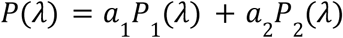

where *a*_1_ , *a*_2_ are values between 0 and 1 representing the normalized modulation of the LED maximum driving voltage to achieve the desired light combination.

In the following of the article, we will use S* , M* or R* to refer to the *N* isomerisations/photoreceptor.seconds in S- and M- opsins, and rhodopsin respectively, resulting from the maximum illumination intensity selected for each stimulus.

#### Cross-opsin activation with LED light illumination

Our light sources also activate rhodopsin. From the close overlap we see in **Fig. 1c** between M-opsin and rhodopsin, their isomerisation rates M* and R* are similar. When our green light is turned on, the ratio between M* and R* is: R*/M*=2. Since we cannot activate rhodopsin or M-opsin independently, we will report only the M* value, since we know that R* will vary accordingly. We cannot rule out that the effects on the responses of ganglion cells can be modulated by the rod pathway, but since we will work on the high-mesopic / photopic level, we expect that the contribution to green color responses will be mostly due to M-opsin activation. The isomerisation rate ratio between M* and S* due to our green light is negligible (S*/M*=1.3×10^-6^). We will report in what follows only the value of M* for the green illumination.

Another unavoidable cross-activation appears when using our UV light, since the tail of the absorption curves of M-opsin and rhodopsin extend well into the short-wavelength side of the spectrum (**Fig. 1c**). When our UV light is turned on, the ratio between M* and S* is S*/M*=4.3, and between R* and S* is S*/R* = 1.2. Although there is less than one order of magnitude difference between of S* and M*, they will be very small compared to the values we will elicit with the green light used in our experiments. In what follows, we will only report the value of S* for the UV light.

### 2.5 Checkerboard stimuli

We displayed a random binary checkerboard at 30 Hz to map the receptive fields (RFs) of ganglion cells, consisting of 40 x 40 checks with a check size of 52.5 µm. This size was chosen to have high spatial resolution in the RFs of neurons, in relation to their average size which is above 100 µm. The stimuli contained 20 s long repeated sequences interleaved with 20 s long unrepeated ones. The former are used to test recording stability, and to quantify reliability in ganglion cell responses. We also use them to calculate firing rates and color onset responses, while the unrepeated sequences are used to retrieve the RFs of ganglion cells by calculating a Spike Triggered Average (STA) from their responses. The checkerboard lasted 30 min for a single color and 1h for interleaved two-color recordings.

#### Building STAs

A three dimensional STA (x, y and time) was sampled using 21 time samples (700 ms). The spatial STA we display in the figures is obtained as the 2 dimensional spatial slice at the maximum value. To display the STAs we zoomed in a 20 x 20 checkerboard (1,05 mm x 1,05 mm) centered on the RF of each cell.

#### Fitting RFs

To visualize the RF of each cell with the STA response, we first do a local smoothing of the spatial STA:

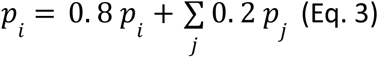

where *p_i_* is any RF pixel coordinate and *p*_j_ refer to the eight neighbor pixels.

Second, we do a normalization so that *max*|*STA*| = 1, and do a thresholding to denoise the background by putting to 0 all absolute values below 0.2. To denoise the signal, we do a second smoothing consisting of an exponential sharpening of the STA with a 1.25 exponent with sign preservation: *sign*(*STA*) · *exp*(*log*(|*STA*| · 1. 25)). The value 1.25 was the lowest that we could find that provided a good fit on the following step. Third, we fit a 2-dimensional Gaussian on the resulting spatial STA. The ellipse corresponding to a contour of 20% from the maximum of the fit is used to delineate the spatial RF. This RF is plotted on top of each spatial STA, when it is detected.

#### Detecting STAs

We evaluated the presence or absence of each cell’s STA at different light levels with a ‘detection’ procedure. We used the RF ellipse contour obtained from the fitting described above and the spatial STA, without smoothing and denoising, and applied the following procedure: first, we check that the center of the fitted ellipse falls sufficiently close to the MEA electrodes position (the central 36% of the full checkerboard area), so that the neuron recorded may have likely received a stimulus input from where the ellipse is located. Second, we calculate the RF area from which we extract an effective radius, as if it was a perfect circle: 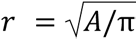. We restrict this radius to lie within 100 and 500 µm as reported in (Baden et al., 2016). Third, we calculate a signal to noise ratio 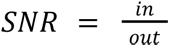, where *in* is the absolute sum of all the STA values inside the RF, and *out* is the sum of the absolute averaged outside of the RF values. Each averaged outside value is obtained by averaging each line of the spatial STA outside of the RF. An arbitrary threshold of 2.75 is applied on this metric which we found was reasonable at detecting an STA. Then, a cell has a detectable STA if it fulfills the three conditions above.

#### Quantifying firing rates

To measure firing rates during the repeated checkerboard sequences, we count the number of spikes in two time windows to separate the transient from the sustained responses. The transient range is 0-2 s and sustained range is 2-20s after a color change in the stimulus. This count is then divided by the number of seconds and the number of repeats to obtain a firing rate in Hz. We used this measure to also quantify the firing rates of the chirp stimulus described below.

#### Quantifying Reliability

To quantify the reliability of ganglion cells responses across repetitions, the average inter-trial correlation was calculated. We considered a 2-20s range from the checkerboard: the unrepeated part of the checkerboard spanned a 20-40 s range, and the 0-2s had a strong transient response due mostly to the previous UV illumination. Therefore, we computed a 2-20s response histogram (bins of 40ms) for each cell and trial, and we calculated the correlation between these histograms, across all possible pairs. The reliability of a cell in response to the checkerboard stimulus is defined as the average of these correlation values. We used this measure to also quantify the reliability of the chirp stimulus described below.

#### Quantifying Recovery

To quantify the recovery during the checkerboard responses, the reliability and the STA detection were calculated on an only green illuminated checkerboard (1 × 10⁴ M*) presented right after the interleaved two-color illuminated checkerboard (1 × 10⁴ M* and 1 x10⁴ S*). This checkerboard was divided in two ways: 1) using 6 consecutive windows of 440 s; 2) using a 1800 s sliding window with a 320 s step size. On these windows, the reliability calculation and STA detection were applied. The ganglion cells were divided into two groups comparing their behavior between M*/S*=20 and M*/S*=1, cells that lost a green STA and cells that maintained a detectable green STA. The values calculated during the green checkerboard during the interleaved two-color stimulus were used as the baseline from which the subsequent recovery was compared.

#### Calculating Recovery time

To calculate the time to reach a steady-state firing rate during the repeated green interleaved checkerboard, we fitted a sigmoid with this formula: 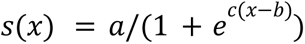 to the 2-40s firing rate histogram (thus ignoring the transient 0-2 s peak). The *τ*, time to return to steady state, was calculated as the time the sigmoid reached 75% of the steady-state value.

### 2.6 Chirp stimuli

We applied a full field (square of 3024 µm side) ‘chirp’ stimulus composed of ON and OFF steps, plus varying frequencies and amplitudes, with luminance values ranging from 0 to 1 (Baden et al., 2016). It was played at 50Hz, containing 20 repetitions of 32 s length each. In the case of the multicolor illuminated chirps we interleaved them, alternating 32 s of UV and 32 s of green illumination. To analyse the responses to the chirp, we created a raster for each neuron by aligning every spike with the chirp intensity profile. Then, we constructed a PSTH (peri-stimulus time-histogram) using a binning of 40ms.

#### Quantifying firing rates and reliability

The same methods were used to quantify the firing rates and reliability of the chirp. These measures were calculated on two parts of the chirp, responses to the steps (0-7s) and responses to the fast modulation (9-27s).

#### Functional clustering

To cluster the ganglion cells we followed the same procedure reported previously for MEA data (Goldin et al., 2022; Gonschorek et al., 2025). To do so, we presented a “white” chirp consisting of a mix of UV and green light at a ratio of M*/S*=20 presented at the same time. From the 217 cells measured in this experiment, we selected 132 cells on the condition to have either a green or a UV spatial STA during a UV/Green interleaved checkerboard at a ratio of M*/S*=20 and whose responses to white chirp had a reliability above 0.1. We used both the responses to the interleaved UV/green checkerboard and the responses to the “white” chirp to cluster the cells in different groups. First, we z-scored the “white” chirp PSTH for all cells and performed a PCA on them. We extracted 16 components that explained above 80% variance of the data. Second, we extracted the temporal profile of the STA of each cell (40 samples at 30Hz) for the two colors. We used the UV STA, and for cells in which it was not detected we used the green temporal STA. We z-scored it and performed a PCA for all cells keeping 1 component explaining 55% of the data. Third, we estimated the spatial STA area by multiplying the major and minor σ values of the 2D fitted ellipse. All values were normalized between 0 and 1. We then built a data vector of 18 values for each cell. We finally performed an agglomerative clustering setting the threshold to have all clusters homogeneous across PSTH and STAs. This resulted in over-clustering producing 24 ganglion cell groups.

#### Pseudo-calcium transformation

We assigned each of these groups to one of the 32 types described in Baden et al., 2016. Due to the different nature of the recordings (calcium imaging and electrophysiology), we first transformed the average PSTH of each ganglion cell group by convolving them with a decaying exponential to match the calcium data (as described in Gonschorek et al., 2025). We then compared pseudo-calcium traces by correlating them with the reference type database (Baden et al., 2016). Finally, we assigned clusters based on these values and merged 22 of our clusters into 16 different ganglion cell types.

### 2.7 Acclimatation stimulus

Before each protocol, we first let the retina adapt to black for 20 minutes and then presented a green (530 nm) color illuminated checkerboard to acclimate the retina during the first 40 to 60 minutes. This green checkerboard was presented at 1 × 10⁴ M*. We adapted only with green checkerboard to prevent adaptation of the ganglion responses to UV, since the lowest UV level projected during our experiments elicits an isomerisation rate of S*= 1 × 10^2^ iso/ph.s. To confirm we were not in a bleaching condition, we did another experiment where we increased M* until witnessing bleaching (**SupFig. 2**). This phenomenon was observed at a much higher value of M*= 1 × 10^6^ iso/ph.s. These acclimatations were done to ensure a reliable response of the retina to our protocol, while waiting also for a proper stabilization of signals and temperature after the retina was placed on the MEA.

**Figure 2.**
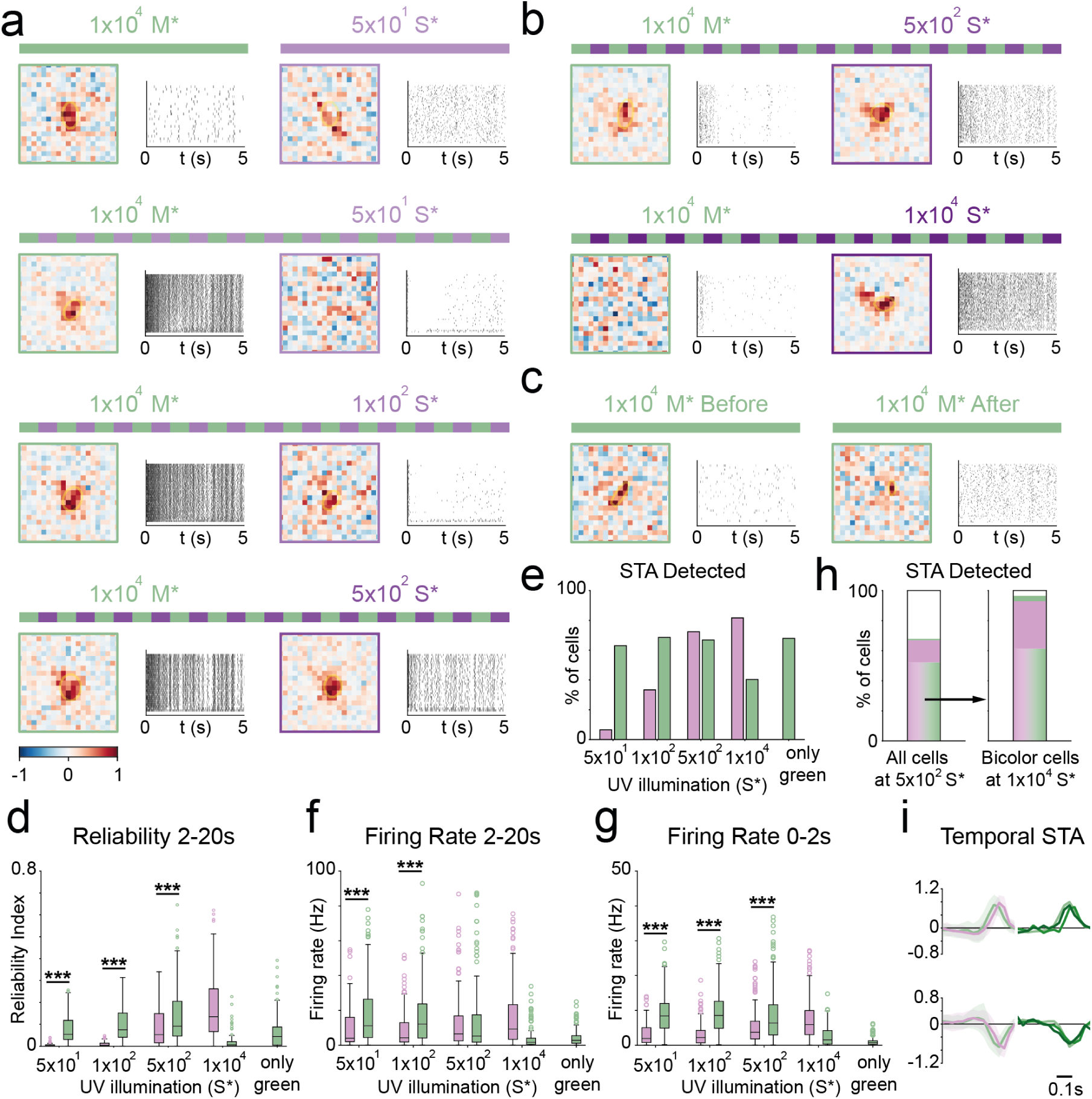
Ganglion cell responses to interleaved UV-green checkerboards with varying UV intensities in the ventral retina. **a-c.** Receptive fields (left) and rasters (right) obtained from the checkerboard stimuli for two example ganglion cells (Cell 1 a, Cell 2 b-c). The first 5 seconds of repeated checkerboards (20 sec) were plotted in the raster. The STAs were extracted with the random checkerboards (20sec) presented after the repeated checkerboards (20sec). Color bar: from response to light output (-1) to response to light input (1). **a.** Example of a cell’s response to interleaved UV-green checkerboards. S* in the UV checkerboard is increased keeping M*=1 × 10^4^ iso/ph.s constant for the green checkerboard, until reaching a close reliability in the raster responses, and a detectable UV STA. **b.** Example of another cell’s response to further increasing S* in UV checkerboard , up to M*/S*=1. **c.** Example of the same cell’s (as in b.) response to only green checkerboard before and after the configurations shown in b. **d.** Reliability of ganglion cell population responses to the paired interleaved UV-green checkerboards tested. (Wilcoxon signed paired test p*<0.001 for green bigger than UV). The last column corresponds to condition c., right: only green checkerboard. The balance configuration lies above 5 × 10^2^ iso/ph.s. **e.** Percentage of STAs detected for the ganglion cells tested with the different interleaved UV-green or only green checkerboards. **f-g**. Firing rate of the population of cells across the paired checkerboards tested. **f.** Sustained response 2-20s. **g.** Transient response 0-2s. The balance pair is slightly below and above 5 × 10^2^ iso/ph.s respectively. **h.** Left: Percentage of ganglion cells with detected STAs for both (violet-green), single (violet or green) or none (white) of the UV-green checkerboards at S*=5 × 10^2^ iso/ph.s. Right: repartition of the cells that had both STAs when the light level changed to S*=1 × 10^4^ iso/ph.s. **i.** ON cells (top) and OFF cells (bottom) population average temporal STAs, normalized. Left: at S*=5 × 10^2^ iso/ph.s and M*=1 × 10^4^ iso/ph.s. Right: green checkerboard STAs with M*=1 × 10^4^ iso/ph.s (light green), M*=1 × 10^5^ iso/ph.s (green) and M*=1 × 10^6^ iso/ph.s (dark green). For d, e, f, g, N = 108 cells for the first two columns, 217 cells for the middle column, and 109 cells for the last 2 columns.

### 2.8 Statistical tests

To compare the differences between the ganglion cell responses to the two interleaved colors, a paired Wilcoxon signed-rank test was used to find when the difference in responses reverted across conditions. This test was used for the reliability and firing rates of the two-color illuminated checkerboards and chirps (**Fig. 2, 5, 6**). These tests were independent one from another and exploratory, hence not requiring any correction to their significance threshold, that was kept at a p-value of 0.05. To test for the recovery of reliability of green checkerboards after presentation of a 1 × 10⁴ S* ultraviolet illumination, a paired Wilcoxon signed-rank test with a Bonferroni correction was used to assess if the reliability increased for each step (**Fig. 3a, 3b**). Finally to test the impact of the cell polarity on firing rates, reliability and firing rates ratios, we used a Wilcoxon-Mann-Whitney test.

**Figure 3.**
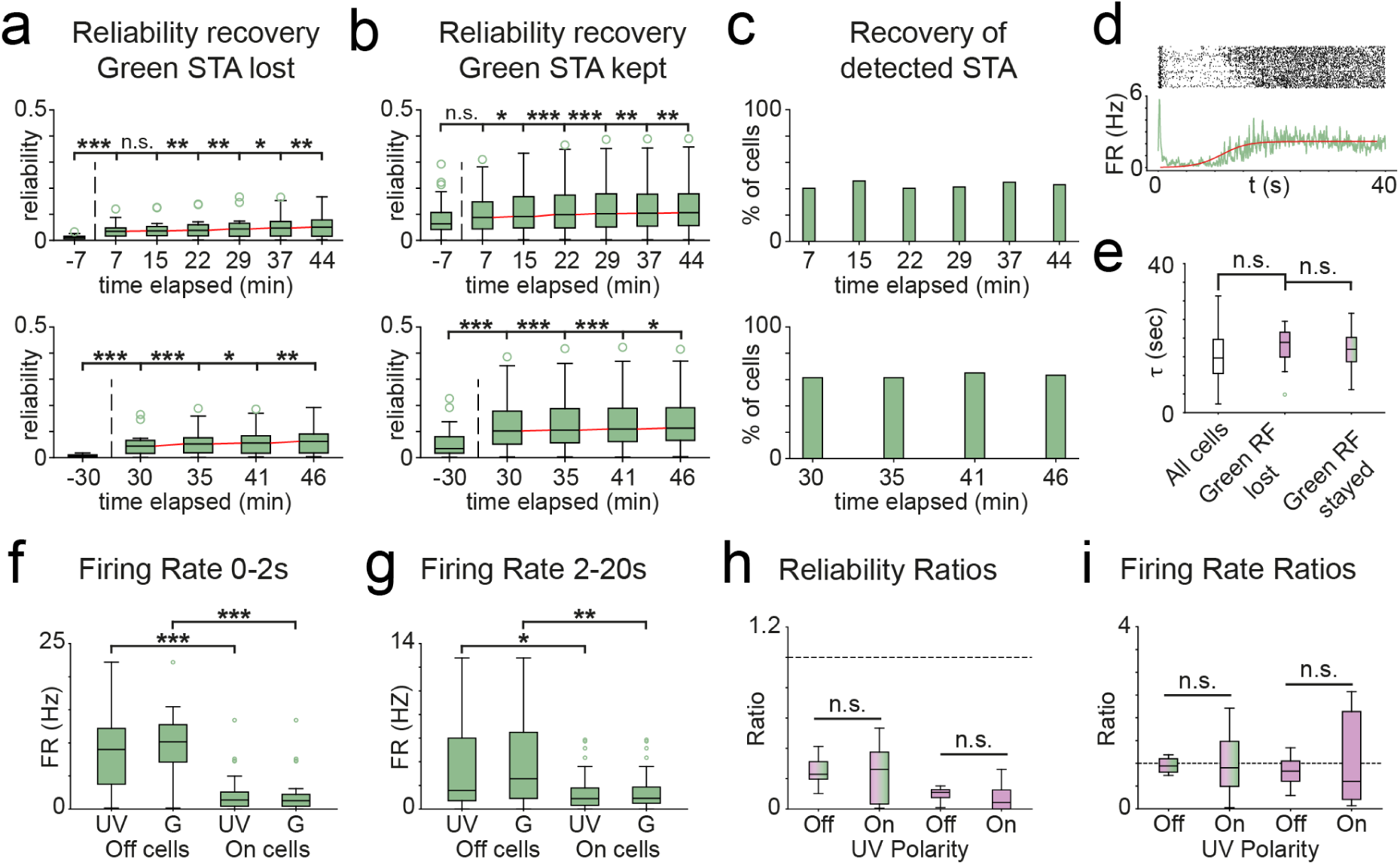
Recovery dynamics of ganglion cell responses to green checkerboard in the ventral retina. **a-b.** Reliability of ganglion cell population responses to the green checkerboard after the interleaved UV-green checkerboard at M*/S*=1. This checkerboard was divided in two ways: 1) using 6 consecutive windows of 440 s; 2) using a 1800 s sliding window with a 320 s step size. (Wilcoxon signed paired test for increase in time with Bonferroni correction, Up: p<0.00016 for ***, p<0.0016 for ** p<0.008 for *, Bottom: p<0.00025 for ***, p<0.0025 for ** p<0.0125 for *) **a.** Reliability recovery for cells that lost their green STA between M*/S*=20 and M*/S*=1 **b.** Reliability recovery for cells that kept their green STA **c.** Detected STA recovery **d.** Example firing rate and histogram for the green checkerboard of a ganglion cell **e.** *τ* population responses shown with box plots, separating ganglion cells with detected and undetected green STAs. Differences between groups is not significant (Mann Withney U test p*<0.3). **f-g.** Firing rate of green checkerboard during the interleaved UV-green checkerboard at M*/S*=1 as a function of green or violet polarity. (Mann Whitney U double-sided test p*<0.001) **f.** Transient firing rate (0-2s) **g.** Sustained firing rate (2-20s) **h.** Reliability ratios responses to the green checkerboard between M*/S*=1 and M*/S*=20 (Mann Whitney U double-sided test p*>0.05) **i.** Firing rate ratios

### 2.9 Daylight spectral intensity curves

#### Single wavelength light

We approximated a light of a single wavelength with a delta function: 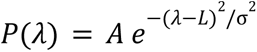, where A is the amplitude, L is the peak wavelength, *λ* is the wavelength, and σ = 0. 01 *nm* is the width we used. We searched for the L values that elicited M*/S* ratios between 10 and 60.

#### Daylight spectra

We used the data from Yu et al., 2023 as natural daylight illumination. We used the data from two sensors, one extracted the spectrum of the light coming from the ground and another recorded the spectrum of the light coming from the sky. The curves were extended to the low wavelengths using the missing data range as reported in Allen et al., 2014. To incorporate the low wavelength data with the correct continuity to the different daytime curves we normalized it, multiplied it by the lowest irradiance value of the different daylight curves, and then incorporated it. Finally, to realistically calculate the isomerisation rate produced by these natural light spectra in the retina of mice, we multiplied them by the transmittance of the full eye of the mouse (Henrikson et al., 2010). The obtained *P_nat_* (*λ*), natural light spectra curves, are then used in equation 2 to obtain S* and M*.

## 3. Results

We will start by showing the disappearance of receptive fields (STAs) from checkerboards with green illumination (from now on green checkerboards), and the absence of high frequency responses from green illuminated chirps (from now on green chirps) when we interleave checkerboards or chirps with UV illumination (from now on interleaved UV-green checkerboards or interleaved UV-green chirps) that have S* in the near photopic range. We will then present the procedure we used to find a range of color balanced S* to M* ratio, matching the reliability of responses across repetitions, and the number of STAs detected under both UV and green checkerboards. We will show that this effect is reversible in the order of a couple of minutes and that it does not depend on single cell firing rates. We will then show that this loss of green STA is not ganglion cell type dependent. In a second phase, we will show that the M*/S* ratio to attain balance is similar for a full field chirp stimulation. We will continue by exploring the range of M*/S* ratio to attain balance along the retinal ventro-dorsal axis. We will end up by estimating the S*-M* ratio elicited in the retina of mice in natural environments and propose a range of illumination that create M*/S* ratios suited to study ganglion cells such that they present reliable responses. We will also quantify ideal narrowband and typical commercial LEDs to see which can stimulate mice photoreceptors within this range.

### 3.1 Equal S* and M* levels prevent ganglion cell responses to green illumination in the ventral retina

We measured STAs in the ventral retina of the mouse in the photopic range using our interleaved UV-green checkerboards (see Methods section 2.5). Our chosen level of M* was similar to the one used in many experiments that used broadband light stimulation in the high mesopic/near photopic level, whether using a single white LED or a simultaneous combination of UV and green sources (e.g. Goldin et al., 2022; Baden et al., 2016, among many others). Similar isomerisation rates are used during optical measurement of retinal activity when using indicators for intracellular Ca^2+^ levels, i.e. calcium imaging, as a proxy to the spiking activity of ganglion cells (Franke et al., 2019; Höfling et al., 2024; Korympidou et al., 2024). We could not find any reference recommending the use of a reasonable ratio between M* and S*, therefore, to start with, we made the S* level to be identical to M*, as it is the approximate ratio used in many studies (Rosa et al., 2016; Stabio et al., 2018; Szatko et al., 2020). One study used a mix of green and UV light with a M*/S*= 30 to determine the location and size of RFs. By varying independently UV and green they extracted the RFs contributions from the M and S opsins (Joesch & Meister, 2016). In our work, the S* and M* values were then both set at 1 × 10^4^ iso/ph.s. In many of the ganglion cells we recorded, the STAs produced by the green checkerboard could not be detected (60% of cells), while the ones produced by the UV checkerboard were present in 82% of the cells (**Fig. 1e**).

Our initial attempt to equalize the activation for S- and M-opsin was inadequate to retrieve both UV and green illumination induced STAs during interleaved UV-green checkerboards. We sought then to study whether a similar effect was present during full field color light stimulation. For this, we stimulated the retina using interleaved UV-green chirps (see Methods section 2.6). The responses to the steps part of the chirp were not abolished (**Fig. 1f**, 0-12 s range), but the fine temporal structure of the ganglion cell response to high frequency and amplitude fluctuations vanished (**Fig. 1f**, 12-32 s range) for most of the cells (47%). We can see that the light step responses at the beginning of the chirp for this example ON cell are still present and slightly diminished compared to the responses to UV, however the response to the fast varying part of the chirp is absent. At a population level, the recorded cells’ activity was more consistent in response to UV light than to green light: more STAs were successfully detected and the response to chirp stimuli was more reliable (**Fig. 1g**).

### 3.2 A S*-M* ratio for reliable color responses to interleaved UV-green checkerboards in the ventral retina

In pursuit of nonlinear summation of green and UV flashed full field illumination, a study (Khani & Gollisch, 2021) found multiple UV-green contrast balance points for different cells. Other works focused on color opponency using full field or they stimulated the center/surround of the cell’s RFs with different spectral content (Szatko et al., 2020; Korympindou et al., 2024). These studies evaluated the strength of the responses during the first hundreds of milliseconds after the ON or OFF illumination steps. They also computed responses to interleaved Green/UV temporal flickers presented during longer timescales 120-300 s. Another study used RGC response to UV and Green flashes to estimate the M/S opsin ratio in the retina (Wang, Weick & Demb, 2011). On the contrary, here we want to explore the effect of high spatial contrast in green and UV at a 30-40 s timescale. We will look for a ratio of M* to S* that provides similarly strong and reliable responses for both UV and green checkerboards. This will in principle allow us to retrieve one UV STA and one green STA per cell, enabling the study of color response structure in a finer scale. While this configuration may not necessarily match the exact color balance present in natural conditions, it may be essential for studying more detailed spatial integration properties of the retinal circuit during color stimulation, and as we will see below it falls very close to it.

To find the M* to S* ratio that balances the reliability of responses to UV and green light, we first verified that an only green illumination with M*= 1 × 10^4^ iso/ph.s allowed us to retrieve green STAs (example in **Fig. 2a**, top, left, during acclimatation checkerboard). We could extract STAs despite the low firing rates of some cells thanks to their high signal-to-noise ratio. We then used on the same retina a low value of S*= 5 × 10^1^ iso/ph.s and found we could retrieve UV STAs in some ganglion cells (example in **Fig. 2a**, top right). With these stimuli we were able to detect STAs in 34% and 27% of cells for green and UV checkerboard respectively (see Methods section 2.5). We have this low value for green checkerboard because it was obtained during the acclimatation; and the low amount of detected STAs for UV was a consequence of the S* used, which did not elicit a reliable firing rate in the cells. We then stimulated the retina with interleaved UV-green checkerboards, keeping the value of M* constant and increasing the value of S* from 5 × 10¹ to 1 × 10⁴ iso/ph.s. When we interleaved UV and green checkerboard with the lowest S* (using the M* and S* we tested above for green and UV only checkerboard), we lost 78% of UV STAs, detecting them in only 6% of the cells (example in **Fig. 2a**, second row). The firing rate in response to the UV portion was also affected, in comparison to when we used a UV only checkerboard (example in **Fig. 2a**, first vs second row). When increasing S*, both the reliability and the firing rate of the ganglion cells in response to UV checkerboards increased (example in **Fig. 2a**, UV raster plots, see Methods section 2.5). In this manner, we found a M*/S* ratio where we detected a similar number of UV STAs and green STAs (N=217 cells, 72% for UV 67% for green, example in **Fig. 2a**, bottom), and whose reliability in the response to the repeated sequences was similar (0.09±0.09 vs 0.14±0.12 for UV vs green average population responses). The balanced configuration we found was for M*= 1 × 10^4^ iso/ph.s and S*= 5 × 10^2^ iso/ph.s, resulting in a ratio of M*/S*= 20 (reliability 0.29 for green and 0.17 for violet, example in **Fig. 2a**).

We then continued to increase the S* until it reached the same value as M* (example cell in **Fig. 2b**). We observed the same effect described in the section above: a strong decrease in the detection of STAs obtained with the green checkerboards. We also saw that the reliability and the firing rate of the ganglion cells in response to the repeated green checkerboard diminished strongly (example in **Fig. 2b**, green raster plots). However, in contrast to the UV raster plot responses when UV STA were undetectable (example in **Fig. 2a**, top right), there remained a strong transient response to the onset of the green checkerboard. This can explain why this configuration, which strongly suppresses green responses, is generally used in the literature to study color responses in ganglion cells: if we restrict ourselves to the first hundreds of milliseconds after a green illumination onset, we can measure a strong firing rate response.

We wondered next if the effects we found were reversible, and presented again the initial green only checkerboard to the same retina. We found that the green STAs came back to the same detected proportion as in our balance configuration (68%, example cell in **Fig. 2c**). Then, to determine a balanced M*/S* ratio for the cell population, we tested whether the reliability was higher in response to UV checkerboards than to green ones for the paired illuminations tested (**Fig. 2d**). We found that the reliability of UV responses surpassed the green ones at the ratio of M*/S*= 1 (S*= M*= 1 × 10^4^ iso/ph.s), while it was the opposite when M*/S*=20. Thus a balanced reliability likely lies between these two values. When we looked for the balance ratio in terms of STAs detectability, we saw that with M*/S*= 20, we had a very similar number of STAs for each color stimulation, with a slightly higher number for UV (**Fig. 2e**), meaning that a balance configuration would have appeared with a higher M*/S* ratio (lower S* for our fixed M*). We then studied the firing rate responses to the green checkerboard, separating them into a sustained (2-20 s, **Fig. 2f**) and a transient (0-2 s, **Fig. 2g**) response. We found that for the sustained response, the balance ratio was achieved in the same range as the one we found studying the reliability. However, for the transient response the balance was achieved in a higher range between M*/S*=100 and M*/S*=20 (Wilcoxon signed paired test p*< 0.01 and p*>0.6).

Regarding the undetectability of the STAs at the ratio M*/S*=1, we studied their evolution starting from the balance configuration at M*/S*=20. All the cells that had only a green STA kept it at the higher S* tested, thus the ones that lost their green STA were found only among the cells having both STAs (52% of the cells). From these cells, 31% lost their green STA, 3% lost their violet STA, 3% lost both, and 61% did not show a change (**Fig. 2h**). This result indicates that for a cross-color effect to be present in the ganglion cells, i.e. UV affecting the color response to green, they have to be sensitive to the information coming both from UV and green.

We then studied the temporal properties of the STAs, to find if there was a noticeable contribution to ganglion cell responses from rhodopsin, through R*. We saw that the green temporal STAs obtained at M*/S*=20 were only slightly slower than the UV temporal STAs (t-test p*<0.01, **Fig. 2i**, left). The difference in latency between both on average was 50ms. This was much faster than the peak responses reported from rhodopsin activation (Joesch & Meister, 2016; Nikonov et al., 2006). It has also been shown that rod-mediated responses are bleached by exposing the tissue to a green LED stimulus delivering R*=1.6×10^6^ iso/ph.s for 2 minutes, and that there is no recovery following the bleach, even after up to 1 hour of measurement (Wang, Weick & Demb, 2011; Wang & Kefalov, 2009). So we applied increasing M* levels and measured the green temporal STAs, without significant changes in their latency (**Fig. 2i**, right), thus showing that most of the responses to the green checkerboard we measured came mainly from M-opsin activation.

### 3.3 Reliability and firing rate decrease do not depend on recovery dynamics nor STA polarity

We wanted to understand why we retrieved less green STAs when an interleaved stimulation with high S* was present. The reason could have most likely been a reduced reliability on the responses. But, was the decrease in reliability during the green checkerboard only caused by a decrease in firing rates? To answer this, we performed a recording of an only green checkerboard right after the last color configuration of M*/S*=1. We displayed this only green checkerboard for 45 min.

Both the reliability and the sustained part of the firing rates quantified for the last green checkerboard showed strongly reduced values, compared to the ones calculated at M*/S*=20 (S*= 5 × 10^2^ iso/ph.s), even if they showed a mild recovery with respect to the M*/S*=1 (S*= 1 × 10^4^ iso/ph.s) (**Fig. 2d,f**). On the contrary, the firing rate of the transient response was diminished (**Fig. 2g**), probably from the now absent contribution of an offset response to UV, which was only present during the interleaved stimulation (see ON-OFF cells responses analysis below). However, the number of detected STAs returned to almost the same value found before presenting the interleaved UV-green checkerboard (**Fig. 2e**). To better understand this recovery process, we studied the evolution across time of the reliability and the detected STAs using smaller time windows to capture recovery speed (see 2.5). We observed that, regardless of the method and time window selected, there was a significant increase of the reliability across the 45 min of recording of the last, only green, checkerboard (**Fig. 3a,b)**. This recovery was slow and, as we mentioned above, did not reach the higher levels displayed near a balanced configuration at M*/S*=20 within this time window. However, after a first important increase from interleaved to green only stimulation, the reliability continued to significantly increase across every time window tested. If we extrapolate this recovery using a linear approximation, we expect to attain the reliability found at the M*/S*=20 condition after 58 mins for the ganglion cells where the green STAs stayed (**Fig. 3a**), and 3hs 57 mins for the cells where the green STAs were lost (**Fig. 3b**).

Regarding the STAs detection, using both the fixed small or the sliding and wider time windows, we saw no significant change of the percentage of ganglion cells displaying a green STA (fisher exact test p*>0.5, **Fig. 3c**). This indicates that the recovery of the cells’ ability to produce a detectable STA in response to green checkerboards lied within a time window between 40s and 7 min, i.e. between the length of one sequence of the interleaved UV-green checkerboard, and the smallest time window we could use to compute an STA. Thus, the first increase in reliability recovery (**Fig. 3a**) seemed sufficient to reestablish STAs detectability in ganglion cell responses to the green checkerboard.

Next, we looked if the recovery could be quantified within the 40 s bout of the green interleaved UV-green checkerboard. In some ganglion cells, we observed a strong transitory peak response at the onset of the green checkerboard, when they were interleaved with UV checkerboards (example in **Fig. 3d**, top). After this transient peak, there was a strong decrease of the firing rate, which was present in most cells. Then, for a large fraction of the cells, there was a slow recovery of the firing rate until a steady state was reached. To verify whether the dynamics of this recovery was influencing the detectability of the green STAs, we quantified the recovery rate of each ganglion cell by fitting a sigmoidal curve and retrieving the characteristic time *τ* it took to the cell to reach steady state (see 2.5). We discarded the first two seconds, where there is the transitory peak.

The cells displaying a quantifiable *τ*<40 s (50% of 109 cells) had an average value of 15.1 ±6.9 ms (**Fig. 3d**, bottom). We divided these cells again in two groups, cells that lost their green STA and cells that did not. We did not find any significant difference in the *τ* of the two groups, indicating that this recovery time was not influencing the detectability of the STA. 42% of the ganglion cells didn’t show any change in firing rate (*τ* = 0 s), from which 22% lost the green STA, 46% kept the green STA and 32% didn’t have a green STA. 8% of the ganglion cells showed a *τ*>40 s, meaning no recovery during the 40 s, 11% of them lose their STA, 79% of them didn’t have a green STA, showing again that this recovery timescale was not impacting the reported change in reliability nor the decrease in the number of detected green checkerboard STAs.

Finally, we wondered if the strength of the transient response was influencing the reliability or the firing rate of the transient period, for the M*/S*=1 ratio. This part of the response to the green checkerboard could have two origins: one due to the onset of the green light, and the other due to the offset of the UV light from the previous sequence. To study it, we divided the cells as ON or OFF, depending on the polarity of their STA. ON cells are defined as responding to a light increase and OFF cells to a light decrease. We did this separation since we expected to find a stronger response to the UV illumination offset in OFF cells at the beginning of the green checkerboard. Indeed, they had a significantly higher firing rate compared to ON cells (**Fig. 3f**), regardless of the color of the checkerboard they responded to. This can be explained because in the majority of them (69%) an OFF cell to green light was also OFF to UV light. The transient firing rate showcased the same behavior: the OFF cells firing rates were significantly higher than the ON cells ones. However, the firing rate was significantly lower during the sustained period between 2-20 s than during the transient one (Mann Withney U double-sided test p*<0.02), as expected from the impossibility of any response to a UV offset then. We did not find any significant difference between cells of the same polarity for different colors within the transient or the sustained responses (Mann Withney U double-sided test p>0.3).

To further study if there was any influence of the firing rate of the cells on the reliability and subsequent detectability of their STA, we separated the cells into ON or OFF again, depending on their STA polarity for the UV checkerboard. We computed first the reliability ratio between an unbalanced configuration M*/S*=1 and a more balanced one at M*/S*=20. This ratio was less than 1 for all the cells, and we found no significant differences between ON and OFF cells (**Fig. 3h**, p*>0.2). However, after comparing the decrease in reliability between the cells that lost their green STA and the ones that did not, we saw a significant decrease in the former (p*<0.05). We repeated the same analysis with the firing rate of the sustained response and we found no significant difference across ON-OFF cells and between cells that had or not their STA become undetectable (**Fig. 3i**, p>0.3).

### 3.4 Green receptive field loss is not ganglion cell type-specific

In response to an interleaved UV/green checkerboard at a ratio of M*/S*=1, some cells lost the green STA they previously showed in response to the same checkerboard at a ratio of M*/S*=20. We wondered then if the cells presenting this behavior belonged to a specific type, for which we did another experiment to classify them by their type. We did the same recording in the upper ventral retina with alternated checkerboards presented at a ratio of M*/S*=20 and M*/S*=100. Our “typing” method uses the cell responses to chirps and checkerboards to cluster them into different groups. We then matched these groups with a reference dataset for mice retina (Baden et al., 2016). After displaying the interleaved UV/green checkerboards at a ratio of M*/S*=20, we presented interleaved UV/green checkerboards at a ratio of M*/S*=1. We found that 16.6% of the cells lost their green STA. We then recorded the ganglion cell responses to a “white” chirp at a ratio of M*/S*=20, composed of simultaneous UV and green light illumination, and applied our typing procedure (See Methods section 2.6). We extracted 24 clusters, and we could assign 22 of them to 16 groups described in Baden et al., 2016.

Within the cell types we found, there was no group with all their cells losing their green STA and 69% of them included at least one cell which lost its green STA (**Fig. 4a**). If we exclude the clusters with less than 5 cells, this percentage rises to 83%, with only ON-OFF Jam-B and ON-alpha types not showing this loss in any cell. Across the 11 groups with more than 5 cells, the percentage of cells losing their green STA had a mean of 27%, showing that this behavior is not specific to any cell type in our experimental configuration. We did not find any remarkable difference between the chirp responses of cells that lose their green STAs and the responses of the cells that kept them (**Fig. 4b**, average Pearson correlation of 0.91 for 11 types), with the exception of some enhanced ON or OFF opposite polarity response to the light steps of the chirp (**Fig. 4b**, see second-to-last line in ON-DS sustained group).

**Figure 4.**
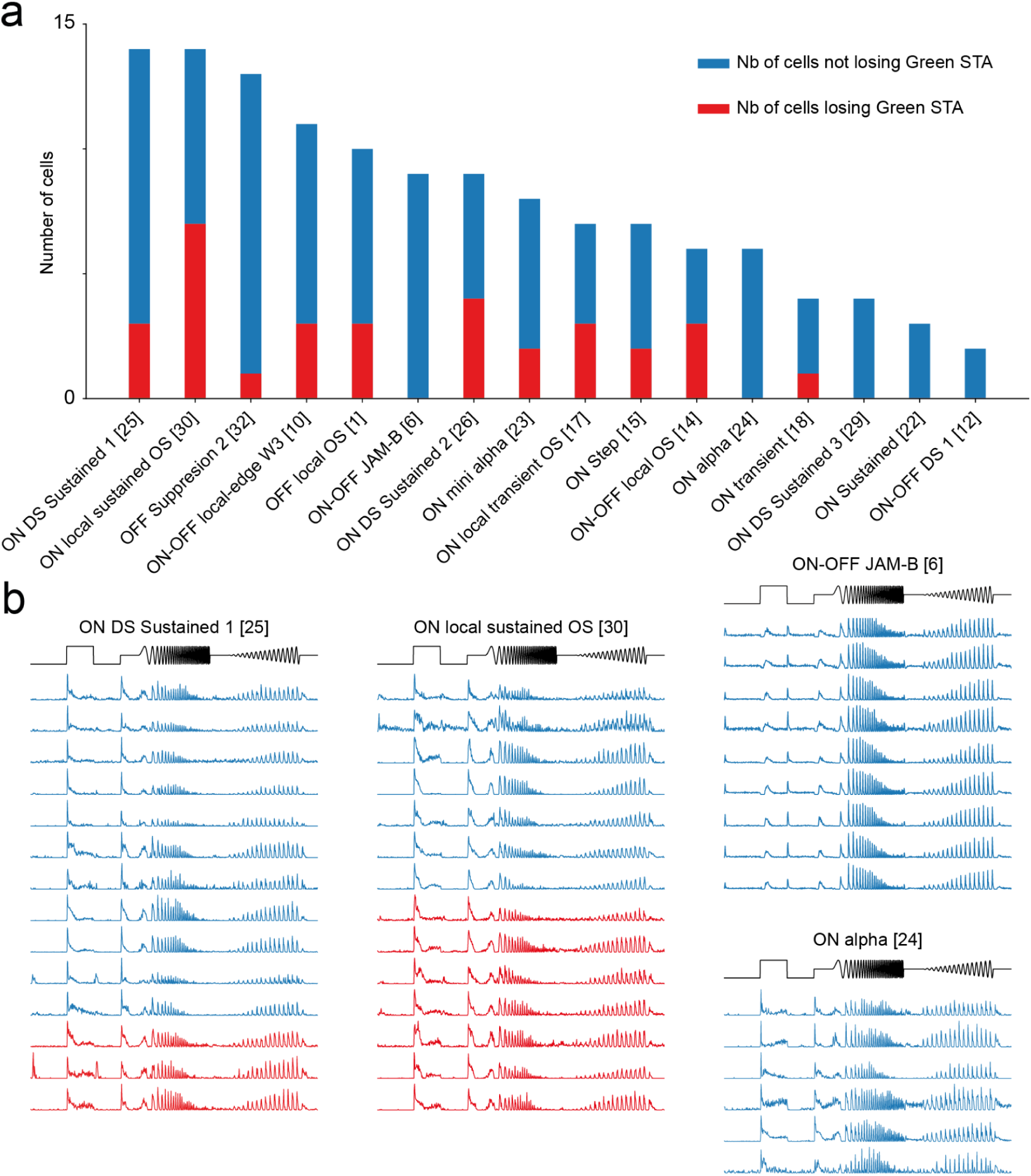
Loss of green receptive fields across retinal ganglion cell types. **a.** Number of cells in each type extracted from one experiment. In red, cells losing their green Receptive Field in response of an interleaved checkerboard passing from a M*/S* ratio of 20 to a M*/S* ratio of 1. In blue, cells keeping their green Receptive Field. **b.** Firing rate traces in response of white chirp for cells in three types (ON DS sustained 1, ON local sustained OS, ON-OFF JAM-B and ON alpha). Cells having their firing traces plotted in red lost their green STA in response of an interleaved checkerboard passing from a M*/S* ratio of 20 to a M*/S* ratio of 1.

**Figure 5.**
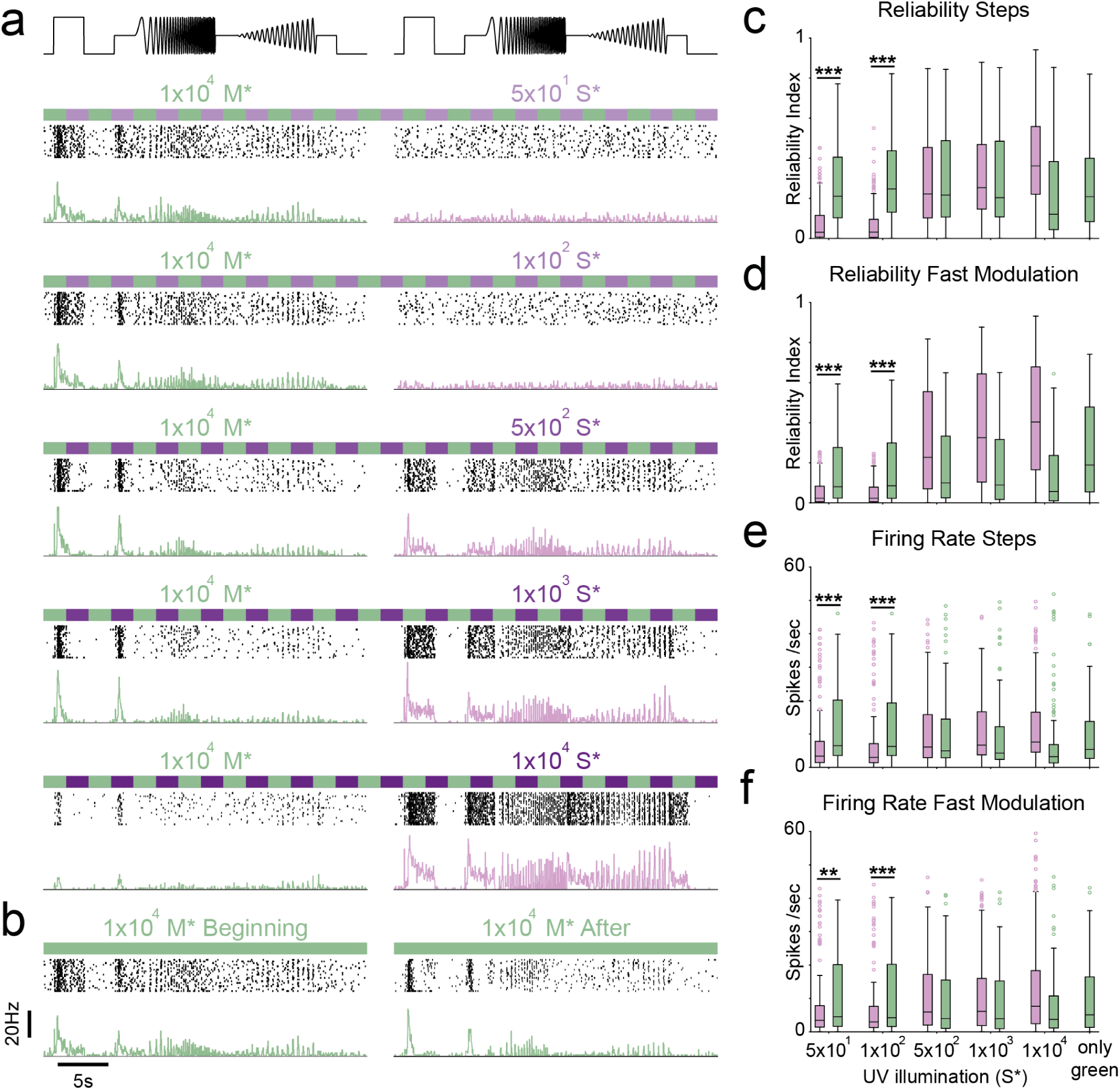
Ganglion cell responses to two-color interleaved chirps with varying UV intensities. **a-b.** Response of an example ON ganglion cell to the interleaved full field color illuminated chirp stimulus. Top: raster, bottom: firing rate. Left: green, Right: violet. **a.** Increasing S* in UV chirps up to M*/S*=1, with constant M*=1 × 10^4^ io/ph.s (149 cells). **b .** Only green chirp comparison in the beginning and after configurations shown in a **c-d.** Reliability of ganglion cell population responses to the paired UV and green interleaved chirps tested. (Wilcoxon signed paired test p*<0.001 for green bigger than UV). Last column corresponds to condition b.: only green checkerboard. **c.** Steps region: 0-7 s. The balance configuration is at 5 × 10^2^ iso/ph.s.(149 cells). Wilcoxon signed paired test (p<0.001 for ***, p<0.05 for *). **d.** Fast modulated region: 9-27 s. The balance configuration is below 5 × 10^2^ iso/ph.s.(149 cells). Wilcoxon signed paired test (p< 0.001 for ***, p<0.05 for *) **e-f**. Firing rate of the population of cells across the paired chirps tested. The balance pair is slightly below 5 × 10^2^ iso/ph.s. **e.** Steps responses 0-7s. **f.** Fast modulation responses 9-27s.

**Figure 6.**
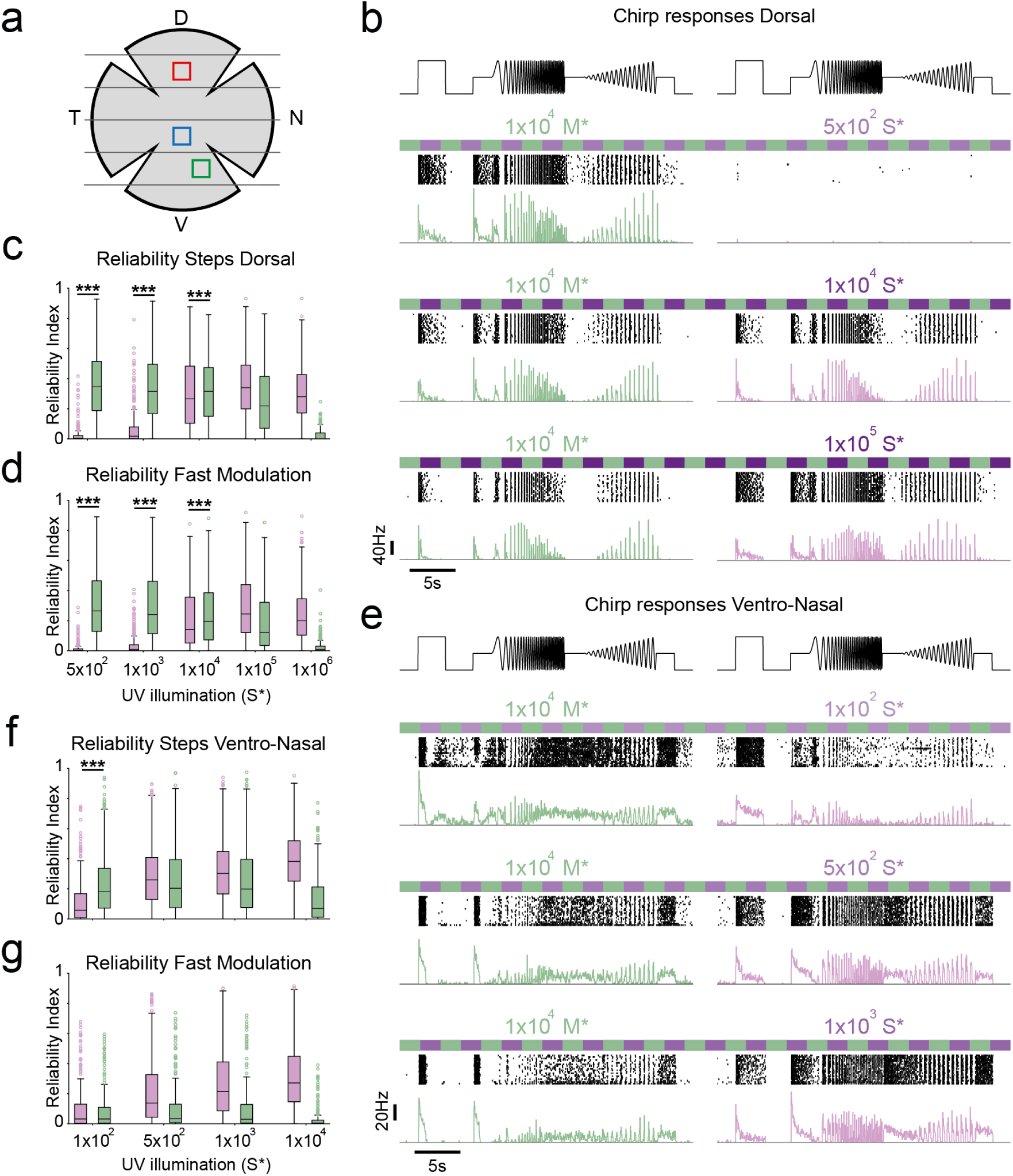
Exploration of the dorso-ventral axis of ganglion cell responses to green and UV. **a**. Location of recorded retinal pieces along the dorso-ventral axis. Blue square: upper ventral retina location, shown in Fig. 2,3,4,5. Red Square: dorsal retina shown in Fig. 6b-d. Green square: lower ventro-nasal retina shown in Fig. 6e-g. **b.** Response of an example ON ganglion cell to the interleaved full field UV/green chirp stimulus in the dorsal retina. Top: raster, bottom: firing rate. Left: green, Right: ultraviolet illumination. Top to bottom: Increasing S* of the UV chirp up to M*/S*=0.1, with constant M*=1 × 10^4^ io/ph.s in green chirps (261 cells). **c-d.** Reliability of ganglion cell population responses to the paired UV and green interleaved chirps tested in the dorsal retina. (Wilcoxon signed paired test p*<0.001 for green bigger than UV). The balance configuration is above S* = 1 × 10^4^ iso/ph.s. (261 cells) **c.** Steps region: 0-7 s. **d.** Fast modulated region: 9-27 s. **e.** Response of an example ON ganglion cell to the interleaved full field color illuminated chirp stimulus in lower ventral retina. Top: raster, bottom: firing rate. Left: green, Right: ultraviolet illumination. Top to bottom: Increasing S* in UV chirps up to M*/S*=1, with constant M*=1 × 10^4^ io/ph.s (257 cells). **f-g.** Reliability of ganglion cell population responses to the paired UV and green interleaved chirps tested in the lower ventral retina. (Wilcoxon signed paired test p*<0.001 for green bigger than UV). **f.** Steps region: 0-7 s. The balance configuration is below S* = 5 × 10^2^ iso/ph.s. (257 cells). **g.** Fast modulated region: 9-27 s. The balance configuration is below S* = 1 × 10^2^ iso/ph.s. (257 cells).

### 3.5 High frequency responses to full field interleaved UV-green color chirps also have an S*-M* optimal balance in the ventral retina

In the previous sections we found a striking reduction in the reliability of the response to green checkerboards when we interleaved UV-green checkerboards with sufficiently high S* in the ventral retina. This produced a decrease in the capability of ganglion cells to reliably respond to fine spatial details, thus preventing the detection of STAs. However, the sustained responses (in the range of 2-20 s from color onset) did not diminish as strongly, and they remained comparable between the cells that had a detectable STA and the cells that did not. We wondered then if a similar behaviour could be found for the finer temporal details in the ganglion cells response to full field interleaved stimulations. For this, we decided to use chirp stimulations, since these are typically used to characterize the responses of ganglion cells and classify them into different cell types (Baden et al., 2016; Gonschorek et al. 2025). This stimulus is 32 s long and contains the typical components used in many studies using color: ON and OFF steps of full field light, allowing us to compare our results with other works.

We can divide the chirp stimulus into two regimes, first 0-7 s corresponding to light ON and OFF steps, and the range between 9-27 s corresponding to a fast modulation of the frequency and the amplitude (see stimulus profile, **Fig. 5a**, top). The protocol applied here was identical to the one we used for the checkerboards, with an extra intensity value at S*=1 × 10^3^ iso/ph.s. Similar to what we described for the color illuminated checkerboards, the green chirp displayed a reliable response that faded away when S* was increased above 5×10² iso/ph.s, reaching very low firing rate values (**Fig. 5a**). The opposite behavior was seen for the responses to UV chirps, as both the reliability and firing rate increased with increasing S*. In the same fashion as with the interleaved green checkerboards, the step responses were not fully abolished, even if for the green chirps these step responses were separated from the previous UV chirp stimulation by a period of dark full field of 4 s. The responses to green stimulation were recovered right after the end of the interleaved UV-green chirp, when we displayed a green-only chirp (**Fig. 5b**). This recovery was much larger for the fast varying portion than for the steps.

When studying the responses to the ON-OFF steps portion of the chirp, we found that their reliability in the UV condition increased with the S*. An equivalence in reliability to the UV and green conditions was found at the same ratio as with the interleaved UV-green checkerboards, i.e. at M*/S*=20 (N= 149 cells, **Fig. 5c**). With the added S* value of 1 × 10^3^ iso/ph.s, corresponding to M*/S* = 10, we determined a higher threshold for the balance configuration, since the green chirp response reliability already showed a decrease. When quantifying the reliability to the fast modulation portion of the chirp, we saw that the balanced ratio appeared for values below S*=5 × 10^2^ iso/ph.s, i.e. for higher values than M*/S*= 20 (**Fig. 5d**). For both portions of the chirp we saw a recovery of the reliability when stimulating with an only green chirp right afterwards (**Fig. 5c-d**, last column).

Regarding the firing rates of the ganglion cells to both the step and the fast modulation regions of the chirps, we saw that they stopped being significantly lower in response to UV chirps than to green chirps when the M*/S*=20 configuration was reached, as S* increased (**Fig. 5e-f**). In both cases, at this M*/S* ratio, we can see that the difference in firing rate between the response to green and the response to UV is not big. These results showed similar behavior in the ganglion cell responses to interleaved UV-green chirps and checkerboards, highlighting the interaction between the two colors on ganglion cell responses, with S* having a strong impact on the reliability of the responses to green color, both for fine spatial and fast temporal features.

### 3.6 Balanced responses of ganglion cells to full field interleaved UV-green color chirps along the ventro-dorsal retinal axis

After studying the responses of ganglion cells to small spatial scales and high frequency modulations of interleaved UV-green stimuli in the ventral retina, we wondered if the ratio of isomerisations M*/S* found to balance ganglion cell responses remained the same across the ventro-dorsal axis of the retina. To answer this, we carried out two dedicated experiments with a similar protocol, one in the dorsal retina and one in the ventro-nasal retina (more ventral than previous experiments , **Fig. 6a**, referred to as “lower ventral retina” from this point on) where the ratio of co-expression of S-opsin in M/S cones is the highest (Nadal-Nicolas et al., 2020).

In the dorsal piece of retina studied, we interleaved UV and green chirps, with the green light producing a constant isomerisation of M*=1 × 10^4^ iso/ph.s, and with the UV light increasing from S*=5 × 10^2^ iso/ph.s to S*=1 × 10^6^ iso/ph.s. Most cells responded only to green at M*/S*=20 (**Fig. 6b**, top row). Increasing UV to reach a ratio of M*/S*=1 made these cells respond to UV and green in a similar way (**Fig. 6b**, middle row). After further increasing the UV intensity to reach a ratio of M*/S*=0.1, the responses of these cells to green decreased slightly but retained their reliability (**Fig. 6b**, bottom row).

Since our UV light also produces a non-negligible M* at high intensities, we wondered if the recorded responses to UV light were in fact just due to M-opsin isomerisation. However, our stimulus was not adapted to answer this question. Nevertheless, we looked across all the 261 cells recorded, and found by eye inspection 11 cells (4.2%, **Sup Fig. 1** for an example cell), whose response to Green was clearly different from their response to UV. Thanks to the differences we saw, in particular to the different polarities found in the responses to green and UV, we could confirm that some cells recorded were responding to S* and M*.

We then repeated, in the dorsal retina, the chirp analysis we did in the upper ventral retina for the ON and OFF steps and the fast frequency and amplitude modulations ranges. For all the cells, the behavior was the same for the steps and for the fast modulation parts of the chirp (**Fig. 6c-d**, 261 cells). As the intensity of the UV LED increased, the reliability of the UV responses increased and the reliability of the green response dropped when S* is above 1×10^4^ iso/ph.s. However, the balanced responses illumination ratio was obtained at a lower M*/S* (0.1<M*/S*<1).

In the lower ventral retina, we performed the same protocol we did before using interleaved UV-Green chirps. We used the same green light level and allowed the UV light to increase from S*=1 × 10^2^ iso/ph.s to S*=1 × 10^4^ iso/ph.s. For most cells, the responses at a ratio of M*/S*=100 were similar in reliability for green and UV, even though the firing rate in response to green was higher (**Fig. 6e**, top row). When we increased UV light to a ratio of M*/S*=20, the firing rate and reliability of the cells’ responses to the fast modulation part of the green chirp decreased compared to their responses to UV (**Fig. 6e**, middle row). And finally, at a ratio of M*/S*= 10, the firing rates and reliability of the responses to green continued to decrease (**Fig. 6e**, bottom row).

If we look at the reliability in the lower ventral retina of the whole cell population (257 cells) in the two regimes described above (the fast modulation and the steps), the reliability behavior was the same as in the ventral retina (**Fig. 6f-g**). The reliability of the responses to UV increased when we increased S*, while the reliability of the responses to green dropped when S* is above 5×10^2^ iso/ph.s. Contrary to what we saw in our previous measurement in the ventral retina piece, the green reliability was low compared to the UV reliability. For the steps responses, the balance in reliability was attained just above a ratio M*/S*=20, similar to the ratio obtained in the ventral retina. However, for the fast modulation response the reliability balance was attained just above a ratio M*/S*=100. Therefore, to attain a balance in lower ventral retina, the ratio M*/S* needs to be between 20 and 100, which is higher than the range of values obtained in the upper ventral retina (60 < M*/S* < 10). This shows that depending on the position of the recording sites in the ventral retina, this ratio can change.

### 3.7 The natural S*-M* ratio of light arriving to mice retina lies close to the ratio that balances UV/green ganglion cell responses in the ventral retina

So far, we studied the responses of retinal ganglion cells to interleaved color stimulations with fine spatial details using checkerboards, and to fast full field modulations in intensity using chirps. The light intensity values we tested indicated that M*/S* = 20 was close to where the reliability to both UV and green stimulations was balanced in the upper ventral retina, for the 40 and 32 s interleaved bouts of color stimuli tested. This value also allowed for a balance in the number of STAs detected during checkerboard stimulation to be reached. Concerning the firing rates, we found that the balance was achieved between the ratios M*/S*=100 and M*/S*=20. We thus decided to use a mid value of M*/S*=60 as a higher cut for finding balanced responses. For the lower cut, we chose a value of M*/S*=10, which we tested for chirps and was close to eliciting a balance too in this part of the retina.

We arrive to an approximate range of 10<M*/S*<60 that supports a balanced reliability in the upper ventral retina and similar firing rate responses to both the UV and green portions of the stimuli (see Discussion for rhodopsin). With this in mind, we wondered if these ratios could be achieved using a unique light wavelength, as it could be the case in many laboratory experimental settings (see Methods section 2.9). We see that a single source of narrow monochromatic light with a peak wavelength between 431 *nm* < *L* < 444 *nm* achieves a balanced-inducing illumination (**Fig. 7a**). However, labs have access to very different kinds of broadly available LEDs. These lights are usually called “white” and are standardized to have an equivalent “temperature” (Kokka et al., 2018). They present a broad spectrum and often a major peak in both a low and a high wavelength ranges (**Fig. 7a**).

**Figure 7.**
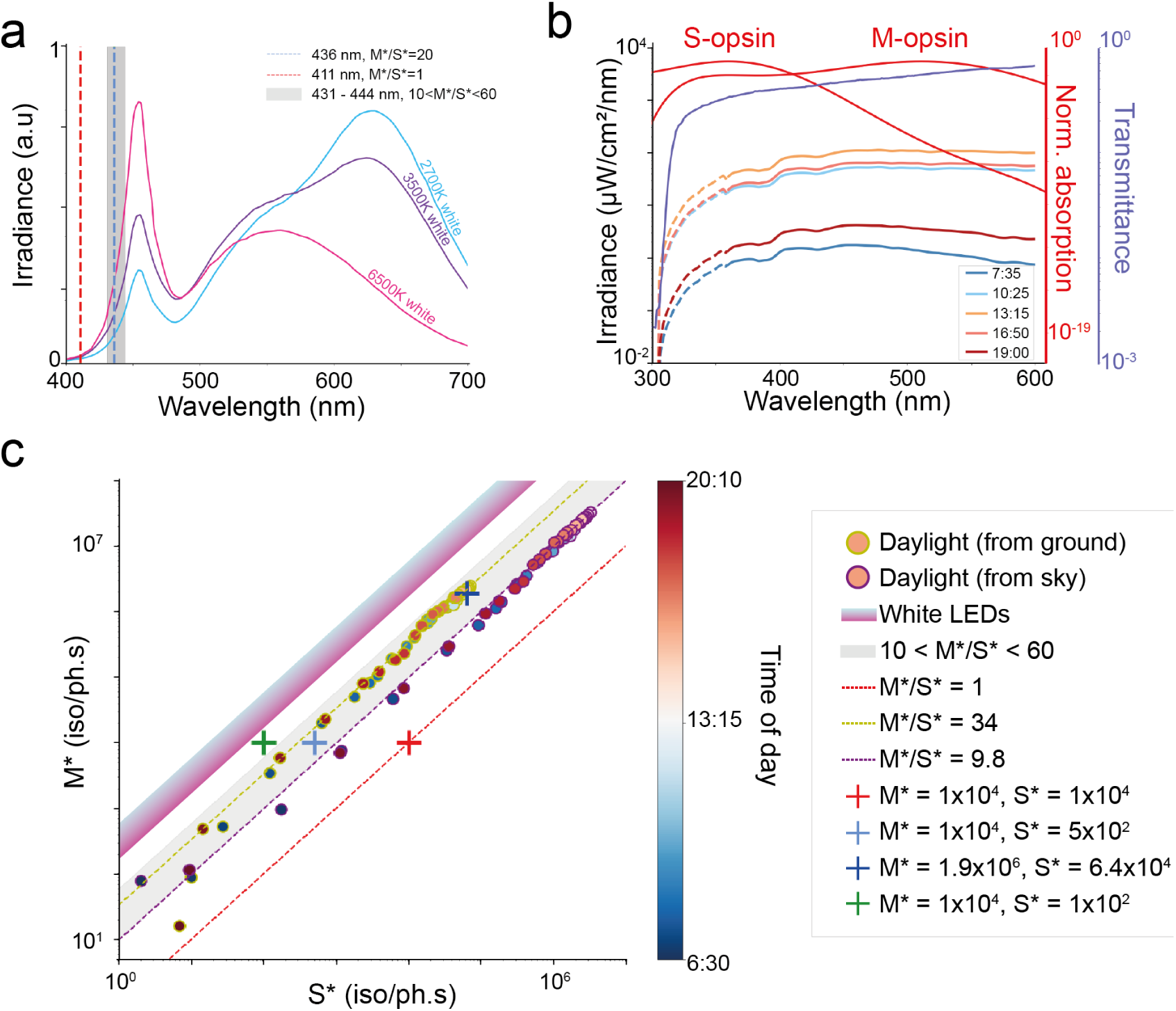
A M*-S* isomerisation rate ratio balancing ganglion cell responses in the ventral retina is close to the natural daylight illumination ratio. **a.** Ideal monochromatic light sources (σ = 0. 01 *nm*) that produce: i) an M*/S*=1 ratio (red) at a peak wavelength of 411 nm, ii) an M*/S*=20 ratio (dashed blue line) at a peak wavelength of 436 nm. The region where these ideal sources produce 10<M*/S*<60 (grey) have peak wavelengths between 431 nm and 444 nm. The normalized irradiance spectra of commercial white LEDS are shown for low (cyan), medium (violet) and high (pink) temperature illumination. **b.** Spectral transmittance of the ocular media of mice before light reaches the retina (purple) together with the measured spectra of natural light at different times of the day (blue to red curves, Henrikson et al., 2010) are overlaid with the absorption curves of S- and M-opsins. The natural light’s spectra have been extended towards short wavelengths (dashed part of the curves, see Methods section 2.9). **c.** M* vs S* iso/ph.s representations of balanced responses configurations found in the retina (blue crosses : upper ventral, green cross : lower ventral, red cross : dorsal retina). The full disks represent the M*/S* ratios of natural daylight at different timepoints (yellow contour: daylight coming from the ground, violet contour: daylight coming from the sky). The balance 10<M*/S*<60 ratio region in the ventral retina is shaded. Dashed lines represent constant M*/S* ratios. White LEDs’ M*/S* ratios are within the cyan-to-pink gradient shaded area, having temperatures ranging from 2400 to 6500 K (cyan to pink, Kokka et al., 2018).

The illumination levels and spectra we use in our experiments are artificial and mice may not find them in natural environments. We quantified then the M*/S* ratio that mice would be confronted to outside of controlled laboratory conditions. To calculate this, we analysed the spectral content of light in natural conditions, taking into account two variables. First, the eye ocular media absorption spectrum, that depends on *λ* (**Fig. 7b**, transmittance curve, Henrikson et al., 2010), showing a strong attenuation of the mouse eye’s transmission in UV, which is the case for many other species (Douglas et al., 2014). Second, the natural spectral content that may change across the time of the day (**Fig. 7b** color curves for sky spectra, Yu et al., 2023). The spectra measured were taken at around 1 meter height, with the light sensor facing up or down (“from sky” and “from ground” in **fig. 7c**), and the recordings were done starting after sunrise and ending before sunset, with a clear (“sunny”) sky. Even if this is not the exact situation a mouse will encounter in nature, it is close to the spectra coming from the sky or the ground, which would arrive respectively at the ventral or dorsal part of the retina, from which we recorded in our work.

With these data, we could calculate the S- and M-opsin isomerisation rates that mice would experience in this natural environment at different times of the day (**Fig. 7c**) (see Methods section 2.9). The data showed a linear relationship, giving a fitted M*/S*=9.8 ratio (R^2^ = 0.9) for the light coming from the sky, and M*/S*=34 ratio (R^2^ = 0.9) for the light coming from the ground. These pseudo-natural rates calculated for the mice retina across different times of the day fall close or inside the region of balanced responses in the upper ventral retina (**Fig. 7c**, gray area), while the unbalanced ratio in the ventral retina and the balanced ratio for the dorsal retina (M*/S*=1, red cross in **Fig. 7c**) lie further away from this region.

To verify this alignment was not merely a coincidence, we performed another experiment, but at a much higher photopic level in the upper ventral retina. Starting with a constant value M*=1.9×10^6^ we increased the S* as we did before, until reaching a balanced response configuration. We did this additional experiment only using chirp stimuli. We found that the balanced ratio was M*/S*=29, with S*=6.4×10^4^ iso/ph.s. (**Fig. 7c**, dark blue cross). This point lies within the gray area defined above, and is closer to the natural daylight data than to the M*/S*=1 ratio (**Fig. 7c**, red dashed line) and near the ratio of M*/S*=20. Therefore, the region of balanced ganglion cells responses in the ventral retina that we empirically found with our experiments lies close to the isomerisation rate ratios the mouse retina may encounter during daylight.

Finally, we asked if the LED lights commercially available are useful in a laboratory setting to probe the mice retina mimicking natural illumination conditions. We see that, on the contrary to a ratio of M*/S*=1, these LEDs will elicit a much higher isomerisation rate for M-opsin than for S-opsin, close to the high unbalanced situation we tested here with low S*=50 iso/ph.s and M*=1 × 10^4^ iso/ph.s (**Fig. 7c**, cyan to violet shaded area). These LED lights are thus eliciting much more activity in the ganglion cells coming from M-opsin activation than from S-opsin.

## 4. Discussion

Our experiments revealed a significant cross-talk between the S-opsin (UV) and M-opsin (green) information processing channels in the ventral part of the mouse retina. There, S-opsin activation strongly influences ganglion cell responses to green light, which elicits mainly M-opsin activation. Equal M*/S* ratios during green-UV interleaved light stimulation produce a suppression of ganglion cell responses to green light. This effect is impacting the capability of ganglion cells to respond reliably to small spatial features (**Fig. 2**) or to fast temporal fluctuations (**Fig. 5**). We did our experiments in the near-photopic level of M*=1 × 10^4^ iso/ph.s, and we confirmed that at a higher photopic level of M*=1.8×10^6^ iso/ph.s a ratio of M*/S* that balances ganglion cell responses also falls within the same region we determined (**Fig. 7**). We also showed that this ratio can change considering the position of the recording sites in the retina (**Fig. 4**) with a ratio that is lower (M*/S*=1) in the dorsal and a higher ratio (M*/S*=100) in the lower ventral retina. We showed that the naturally occurring mesopic/photopic daylight illumination conditions reaching the retina of mice from the sky, from dawn to dusk, produces an approximate constant isomerisation rate ratio near M*/S*=10, which lies on the limit of the range we found empirically for the ventral retina. Whereas the ratio produced by natural light coming from the ground is at M*/S*=34, within the confidence range we found in the ventral retina. This finding is in accordance with an information theoretical approach (Barlow, 1961), where it is suggested that to efficiently encode information, sensory relays need to be specifically tuned to the stimulus they want to encode (having a M*/S* encoding ratio near the natural one).

To elicit comparable ganglion cell responses in the ventral retina, a 1 to 2 orders of magnitude for M* to S* is required. This disparity suggests that UV-activated pathways undergo a different amplification mechanism compared to green-activated ones, which is witnessed by a higher sensitivity at shorter wavelengths (Jacobs, Neitz & Deegan, 1991). Previous works have also shown the suppression of responses to green at high isomerisation rates using achromatic light, and argued that this is due to signals coming from the rod pathway which reaches saturation and stops responding (Joesch & Meister, 2016; Nikonov et al., 2006). Although there may be some rod contribution to green light responses, and it has been shown that rods can escape saturation (Adelson, 1982; Khani & Gollisch, 2021), this is not the effect we show here for the 52% of the ganglion cells that show responses to both UV and green illumination. At the ratio of M*/S*=20, our interleaved UV illumination only elicits M*=1.2 × 10^2^ iso/ph.s, and our green light kept M* constant at least two orders of magnitude above, together with R* in close to saturation levels (2 × 10⁴ iso/ph.s).

Two observations suggest that the biggest contribution to the responses we measured in response to green stimuli in the ventral retina come from M-opsin cone activation and not from rods. First, the temporal STAs responses were fast for the green checkerboards, while it is known that they should be much slower for rhodopsin inputs. When stimulating with UV light, activating mostly S-opsin cone pathways, mice ganglion cell responses are typically faster, similarly to those of primate ganglion cells to visible light (Demb et al., 2015; Wang, Weick & Demb, 2011; Borghuis et al., 2014). The responses we recorded had similar latencies for both the UV and green checkerboard, and they did not change even when we increased the luminance to reach bleaching levels for rhodopsin (**Fig. 2h**). Second, we measured only 1 color opponent cell out of 432 cells, looking only at their STA polarity. It has been shown that color opponency in mice is mainly driven by an S-opsin/rhodopsin input coming through lateral feedback from horizontal cells (Joesch & Meister, 2016; Szatko et al., 2020). Horizontal cells can form a dense, electrically coupled network (Baden et al., 2019; Becker et al., 1998; Cook & Becker, 1995) that interacts with photoreceptors via lateral inhibition. In our experiment, only the ganglion cells that displayed only green STAs were unaffected by the interleaved UV light. This suggests a mechanism by which both S-opsin and M-opsin inputs converge at a lower processing stage in the retinal circuit, or happen to be processed from co-expression of S- and M-opsin within the same cone (Szel et al., 1992; Haverkamp et al., 2005).

In the lower ventro-nasal retina, the ratio to attain balanced ganglion cell responses for the light steps of the chirp was in the same range as in the ventral retina (M*/S* between 10 and 60). In contrast, for the fast modulation part of the chirp, a ratio of M*/S*=100 is needed (**Fig. 6d**). This indicates that this ratio is stimulus dependent. In addition, depending on the region of the ventral retina recorded, this M*/S* ratio can vary in a wide range (from 20 to 100 in response to fast modulations). This is in line with the fact that the S-opsin expression in the ventral retina is not constant, and these differences may explain the regional change in the ratios we found. But, is there a relation between the M*/S* ratio we found that balances ganglion cell responses across the ventro-dorsal retinal axis and the actual M/S density ratio of opsins? In Wang, Weick & Demb, 2011 they patched cells along the ventro-dorsal axis to find the balance point where UV and green responses to flashes were similar by changing the ratio of M*/S* similarly as we do here, but with simultaneous color stimulation. From this, they calculated that the ratio of S/M opsins lies between 134 and 5.5 across the ventral retina. This follows closely the ratios that we found in the upper ventral and lower ventro-nasal retina with M*/S* between 100 and 10. It could be that the reason why the ventral retina seems optimized at balancing ganglion cell responses to a natural light spectrum is because the product M*/S* x M/S may be stable at different ventral retinal locations.

On the contrary, in the dorsal retina, the ratio we found is closer to 1 (**Fig. 6e-g**). This may reflect the low density of S-opsin in the dorsal retina, which was reported to be of only 4% (Nadal-Nicolás et al., 2020). However, we cannot affirm that the decrease in reliability for the green responses in the ventral retina and the dorsal retina when increasing UV illumination is due to the same phenomenon. In the dorsal retina, when we present interleaved light with a M*/S* ratio below 1, our UV light also produces M*=2.3 × 10^4^ iso/ph.s, while eliciting S*=1 × 10^5^ iso/ph.s (See methods section 2.4). The reliability decrease of the green responses could be due to the impact of S-opsin isomerisations, as in the ventral retina, or it could be due to the adaptation of the retina to the slightly higher M* produced by the UV light. When the retina adapts to this higher M*, it may not respond with as much reliability and firing rates to the lower M* produced by the green light. Therefore, we saw an interaction coming from the UV on the green light responses, but we can’t conclude if it is due to the adaptation of the retina to the M isomerisation of the UV light or the S isomerisation as in the ventral retina.

When analyzing the group of ganglion cells that lost their green receptive fields in the upper ventral retina due to UV light stimulation during the interleaved color checkerboard, we found that they did not belong to any specific cell-type, as defined in Baden et al., 2016. We found only two types (with more than 5 cells) that did not have cells that lost their green STA: the ON Alpha and the ON-OFF JAM-B. Further recordings are needed to verify if these specific cell types don’t have a cross-talk effect between UV and green.

Different dynamics of recovery are found in the retina, and we have also found recovery of ganglion cells responses at different timescales. First, during an interleaved green-UV checkerboard at a ratio of M*/S* = 1 , we saw in some cells that the firing rate in response to green stops and recovers in 10-40 s. Second, after the end of this interleaved checkerboard, the detected STA recovers in minutes (1-7 min) but the reliability takes longer to recover (1-4 h). In the same way, photoreceptors are capable of reacting at multiple timescales. Fast dynamics were found when modeling photoreceptors adaptation mechanisms (1-10 ms, Angueyra et al., 2022; Idrees et al., 2024). These photoreceptor models, which capture key biophysical mechanisms, improved predictions of retinal responses to natural stimuli across wide isomerisation rate changes. On the other extreme, dark adaptation takes 3 hours to span 5 orders of magnitude in light sensitivity in cats (Barlow et al., 1957). Another slow adaptation is the recovery of sensitivity to bright light after dim light exposure, due to rapid photopigment breakdown, causing bleaching and glare (Cornwall et al., 1990). These two types of adaptation were not present in our study. Our detected STA recovery time seems compatible with the timescale reported in human psychophysics experiments related to color persistence (Gupta et al., 2020). This effect is called “chromatic adaptation”, and it consists of the human visual system’s capability to adjust to widely varying colors of illuminants in order to approximately maintain the consistency of perceived color appearance. The time course of chromatic adaptation can be divided into 3 stages: an extremely fast, a fast and a slower stage (Fairchild & Reniff 1995; Rinner & Gegenfurtner 2000). It was found that the slow chromatic adaptation reaches 80–90% within a minute and is complete after five minutes. Further work is needed to determine if the timescale we measured here in mice, and the slow adaptation required for the colour constancy to stabilize in humans, are produced by the same mechanisms.

Future studies may explore smaller time intervals in between color interleaved stimulations, to see if our results hold for intervals below 40 s / 32 s. It could be interesting to see if the effect disappears in a continuous manner, if there is a fast transition, or if it remains the same. This can be achieved using two projectors, one for each color, as shown in our work or in this study (Franke et al., 2019). This will allow us to better understand the computations of the ganglion cells, and of the retina in general. Simultaneous two-color light stimulation may reveal on the other hand strong nonlinear interactions, at least for the transient responses lasting up to several hundreds of milliseconds. (Khani & Gollisch, 2021). Therefore, studies using linear decorrelation to extract the contributions of one opsin from the other to the responses could have a bias due to the non-linearity of color processing in the retina.

Regarding the number of responding cells, our work may explain the strong unbalance found in previous works between the number of ganglion cells presenting a center RF for UV light compared to green light in the ventral retina (Szatko et al., 2020). It could also explain why there are more amacrine cell classes found tuned to UV compared to green (Korympidou et al., 2020). This effect could be due to the M*/S*=1 isomerisation rate ratio used in these studies. We have witnessed in our work a decrease in 60% of the ganglion cells displaying a green STA when arriving to this ratio, from the balance ratio of M*/S*=20 (**Fig. 2e**). The balanced ratio found in our work highlights the complexity of color processing in the retina and the necessity of fine-tuning stimulus spectral content and intensity to accurately study color interactions in visual processing. Our checkerboard stimulus is tailored to study small spatial details and extract STAs, so it does not invalidate that color opponency in mice is present mostly in the cells of the ventral retina. However it may change the view in which the ventral retina of mice is mostly dedicated to S-opsin signal processing, since M-opsin responses appear widespread through the ganglion cells we recorded.

Previous works used “white” or 415 nm LED to illuminate the mouse retina for different stimuli. We saw here that using these types of illuminations will not create a M*/S* ratio found in natural conditions. On the one hand, a “white” LED will have a M*/S* ratio superior to the natural ratio, causing the retina to process only the green inputs. Using a single color LED that has a M*/S* < 10 will cause the retina to process only the violet inputs in the ventral retina. Thus, a bias towards studying only the cells not abolished by UV light or by green light is likely to be present in previous studies, depending on the type of illumination used. A solution for this is either to calibrate the intensity of two or more light sources to attain a balanced M*/S*, or to buy a single blue light source that achieves this (**Fig. 7a**).

If we want to study color response in the retina, it’s important to study the surround of the receptive fields because the color opponent cells encoding color information work with a center/surround mechanism (Joesch & Meister, 2016; Szatko et al., 2020). However, our checkerboard stimulus was not suited to extract surround responses This is due to the small checks used to obtain with good resolution the center of the receptive field of ganglion cells, which are not big enough to elicit sufficient responses from the surround.

Regarding fast varying temporal responses, we attained the same balanced responses with a natural M*/S* ratio (**Fig 5c**). For the ON and OFF steps some green responses stayed, even at a lower M*/S* ratio. We don’t know if this is enough to study the smaller spatial details of fast varying natural scenes. This information may not be used effectively by mice, or it could be relevant for their natural behavior in natural illumination conditions. Recent works have explored the retina and V1 responses to color movies (Franke et al., 2022). These movies were taken with a novel camera that measured precisely the spectral content of the natural habitat of mice (Qiu et al., 2021). From these videos, they extracted the ratio between UV and green intensities and therefore the ratio of M*/S* created by some images. They calculated that the ventral retina receives a M*/S* ratio between 1.4 and 1.6 and that the dorsal retina receives a M*/S* ratio between 4.5 and 6.5. These results are coherent with ours regarding the bigger values arriving in the ventral with respect to the dorsal (**Fig. 7c**), they differ however by a factor of around 7. These differences may be due to the different positions of the camera in the two studies, on the ground vs 1m above ground in our case. They could also be influenced by the different sensors used: a camera with a filter, vs an spectrometer that captures the whole light spectrum in our case. Moreover, the difference between their ratios and ours can also be due to them reporting average isomerisation rates across an image, while we indicate maximum nominal isomerisation rates. From our findings, we show that we may need to adjust the UV/green illumination to avoid an unnatural shift of the responses to only the UV illumination channel. When considering the ratio we obtained at a high mesopic/photopic regime in the ventral retina, it could seem unnatural for mice which are typically regarded as strictly nocturnal. However they often forage during the day in the wild (Daan et al., 2011). Thus, studying the natural illumination from dawn to dusk, from mesopic to photopic may be useful to understand the way mice use their S- and M-cones in a natural setting.

In conclusion, our findings highlight the need for a precise color illumination balance in photopic conditions when studying small spatial and fast temporal color response features in ganglion cells. This will place the ventral retina, contrary to the dorsal one, into a close to naturally occurring illumination condition, which allows the M- and S-opsin information to arrive simultaneously, and with the same reliability, to the majority of the ganglion cells. However, in the dorsal retina it would be more suited to use a M*/S*=1 ratio, from the scarcity of S-opsins, relative to M-opsin. In future studies that explore the responses to color natural images or videos, maintaining a proper color balance will be essential. It will help disambiguate from an instantaneous nonlinear color interaction, when two colors are presented simultaneously, from the long lasting effects we show here, ensuring a more accurate interpretation of ganglion cell responses, and retinal processing in general. Choosing appropriately the stimulus intensities will help avoid misinterpreting color interactions in the ventral retina visual processing. This has broader implications not only for retinal studies but also for understanding color vision in other species and in the context of color vision restoration efforts (Bansal et al., 2024). Future research should consider these factors when designing experiments and interpreting data, particularly when exploring visual processing of complex, dynamic color scenes.

## Supplementary Figures

**Supplementary Figure 1.**
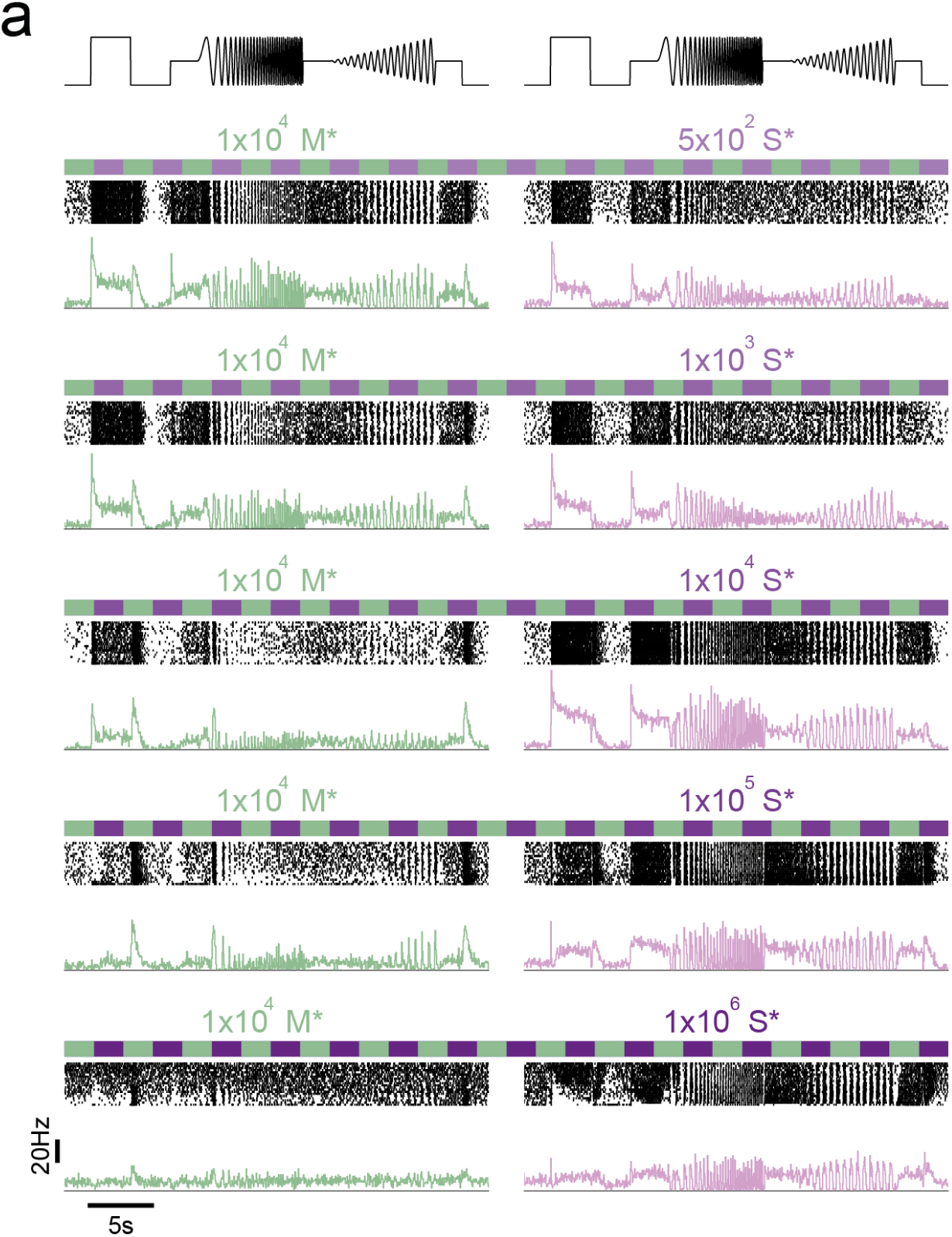
Example of a cell responding to UV stimulus in the dorsal retina. **a.** Response of an example ON ganglion cell to the interleaved full field color illuminated chirp stimulus in dorsal retina. Top: raster, bottom: firing rate. Left: green, Right: violet. Increasing S* in UV chirps from M*/S*=20 up to M*/S*=0.01, with constant M*=1 × 10^4^ io/ph.s. We see that only green illumination elicits an OFF response for all UV intensities tested.

**Supplementary Figure 2.**
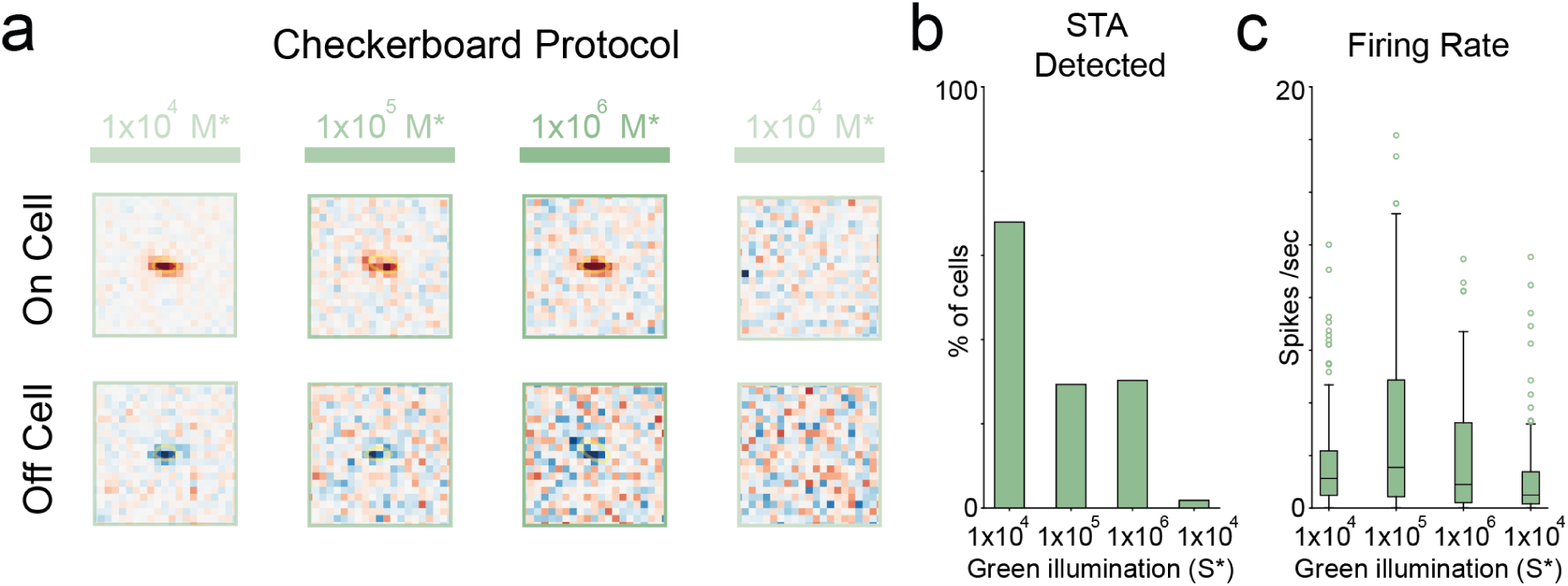
Bleaching the retina in response to high M*. **a.** Receptive fields of an example ON ganglion cell and an example OFF ganglion cell in the ventral retina. We extracted an STA from their response to green checkerboards stimuli (30 min each), increasing M* from 1 × 10^4^ io/ph.s to 1 × 10^6^ io/ph.s and coming back to 1 × 10^4^ io/ph.s. **b.** Percentage of STAs detected for the ganglion cells tested with the different green checkerboards. **c.** Firing rates of ganglion cell population responses to the green checkerboards tested.

## Data statement

Data will be made available upon publication.

## Acknowledgements

We would like to thank Olivier Marre for helpful discussions. This work was funded by Agence Nationale de la Recherche grants ANR-23-CE37-0004-01 HiDeepID (MG), ANR-22-CE37-0016-01 PerBaCo (MG), ANR-25-CE19-3927-02 REFOCUS (MG), ANR-21-CE37-0024 NatNetNoise (UF), by IHU FOReSIGHT ANR-18-IAHU-01 (UF), by Sorbonne Alliance Emergence SHARPEN (MG), and by Sorbonne Center for Artificial Intelligence-Sorbonne UniversityIDEX SUPER 11-IDEX-0004 (UF). This work has been done within the framework of the PostGenAI@Paris project and it has benefitted from financial support by the Agence Nationale de la Recherche (ANR) with the reference ANR-23-IACL-0007. Our lab is part of the DIM C-BRAINS, funded by the Conseil Régional d’Ile-de-France.

## Author contributions

TQ, AL, FC and RB performed experiments. TQ, AL, FP, UF and MG analyzed the data. TQ, AL, MG wrote the manuscript with input from all authors. MG and UF conceptualized the work. MG supervised the work. MG and UF secured funding.

## Notes

### Competing Interest Statement

The authors have declared no competing interest.

### Summary of Updates

We performed additional experiments, analyses and rewriting of the manuscript. In particular, we explored more retinal regions, dorsal and ventro-nasal, showing that the balancing ratio varies across the retina, and that the ventral retina is the region that aligns better to the natural light spectrum. We typed the cells of a new experiment in the ventral retina and found that there is no specific type that shows a loss of green responses due to UV light. We changed the title, some wording of the summary, and the end of the introduction to reflect our new findings. We added two figures and two supplementary figures to reflect the new data and the new analyses made.

